# Towards understanding predictability in ecology: A forest gap model case study

**DOI:** 10.1101/2020.05.05.079871

**Authors:** Ann Raiho, Michael Dietze, Andria Dawson, Christine R. Rollinson, John Tipton, Jason McLachlan

## Abstract

Underestimation of uncertainty in ecology runs the risk of producing precise, but inaccurate predictions. Most predictions from ecological models account for only a subset of the various components of uncertainty, making it difficult to determine which uncertainties drive inaccurate predictions. To address this issue, we leveraged the forecast-analysis cycle and created a new state data assimilation algorithm that accommodates non-normal datasets and incorporates a commonly left-out uncertainty, process error covariance. We evaluated this novel algorithm with a case study where we assimilated 50 years of tree-ring-estimated aboveground biomass data into a forest gap model. To test assumptions about which uncertainties dominate forecasts of forest community and carbon dynamics, we partitioned hindcast variance into five uncertainty components. Contrary to the assumption that demographic stochasticity dominates forest gap dynamics, we found that demographic stochasticity alone massively underestimated forecast uncertainty (0.09% of the total uncertainty) and resulted in overconfident, biased model predictions. Similarly, despite decades of reliance on unconstrained “spin-ups” to initialize models, initial condition uncertainty declined very little over the forecast period and constraining initial conditions with data led to large increases in prediction accuracy. Process uncertainty, which up until now had been difficult to estimate in mechanistic ecosystem model projections, dominated the prediction uncertainty over the forecast time period (49.1%), followed by meteorological uncertainty (32.5%). Parameter uncertainty, a recent focus of the modeling community, contributed 18.3%. These findings call into question our conventional wisdom about how to improve forest community and carbon cycle projections. This foundation can be used to test long standing modeling assumptions across fields in global change biology and specifically challenges the conventional wisdom regarding which aspects dominate uncertainty in the forest gap models.

## 1 Introduction

Understanding predictability is an ecological grand challenge because ecological predictions provide both a road map for scientific learning and a practical tool for real-world decision making. One of the key ways to measure predictability is to estimate uncertainties in predictions and how they grow/decline (Dietze, 2017b). The overall uncertainty can be partitioned into variance components allowing us insight into which aspects of the modeling process contribute to the accuracy and precision of an ecological prediction. In models of community and population ecology, the variance components of overall prediction uncertainty are: demographic stochasticity, internal state (i.e., initial conditions), external forcing (i.e., drivers/covariates), parameters, and modeled processes (Box 1).

Quantification of uncertainty in global change ecology studies have traditionally focused on demographic stochasticity, parameter uncertainty, and meteorological forcing uncertainty. The inclusion of demographic stochasticity through variation in demographic rates among individuals adds realism to population predictions and has been theoretically proposed to be important for explaining species coexistence (Tilman, 2004). However, it has not been shown that this component of uncertainty is especially important in other aspects of model predictability, like prediction of abundance or mass. Second, uncertainty in external forcings, such as climate drivers, are often incorporated into forecasts because of large uncertainties about future environmental states that are dependent on scenarios (anthropogenic emissions, land use, etc.) about human decisions and behaviors (Bonan, 2015). However, over shorter timescales that are less sensitive to human scenarios (e.g. daily through decadal) there can still be considerable uncertainty about environmental drivers. Even in retrospective “hindcasts”, acknowledging uncertainty in past external environmental drivers is important for accurately attributing causal relationships between drivers and resulting ecosystem states. Lastly, parameter uncertainty has become a dominant focus of calibration and uncertainty studies (Fischer et al., 2019; Fisher et al., 2019; Fer et al., 2018; Reichstein et al., 2019; Raczka et al., 2018). Process models in particular often have large numbers of parameters that historically have been under-constrained by data. Even when constrained by data, model parameters have often been optimized to a single “best” estimate that ignores both the real uncertainty in parameter values and the common tendency for parameters to trade-off or co-vary with one another. Constraining both external drivers and parameters with data has greatly improved process-model performance and shown that as data volumes increase parameter uncertainty tends to decline asymptotically (Fer et al., 2018; Dietze, 2017b).

In contrast with the attention given to demographic stochasticity, environmental driver scenarios, and parameters, the uncertainty from initial conditions and process uncertainty are seldom considered in ecological forecasts. On the one hand, ecological systems are less sensitive to initial conditions than deterministically chaotic meteorological systems (Lorenz, 1963; Rabier et al., 1996), and some studies have found initial conditions uncertainties are often small and decay quickly with time (Bonan et al., 2019; Cox and Stephenson, 2007). On the other hand, there is substantial historical dependence in ecology (Ricklefs, 1987), and many important ecological processes have slow dynamics and long memory (e.g., forest succession). It seems prudent to consider the impacts of initial condition uncertainty. Pragmatically, initial condition uncertainty was often omitted from ecological forecasts because appropriate data to constrain the variety of initial states in complex ecological models were rare. Fortunately, increasing amounts of coordinated, large-scale ecological data are being collected to constrain these uncertainties (remote sensing, inventory and monitoring data, coordinate research networks, etc., LaDeau et al. 2017) that allow us to test how much initial condition uncertainty affects prediction.

Process uncertainty is even less frequently quantified but is also important to include because it represents the uncertainty in prediction caused by model simplifications and assumptions (Wikle, 2003; Clark and Bjørnstad, 2004; Cressie et al., 2009). In principle, process uncertainty can be estimated in retrospective studies by comparing the distribution of modeled state variables to observed state variables. However, this is not as simple as calculating a RMSE between modeled and observed time series. Estimating process uncertainty requires a robust approach for partitioning, at every point in time, the observation errors in the data; the uncertainties about the previous state of the system; and the contributions of parameter uncertainty, driver uncertainty, and demographic stochasticity to the growth in error over that time step. Such an approach has been possible for simple process models within a state-space modeling framework (Clark and Bjørnstad, 2004; Patterson et al., 2008), but has not been available for complex models because the estimation process is too computationally demanding.

To address this issue, we develop a method for fully partitioning the five types of uncertainty (Box 1) in complex process-based ecological models including a novel generalized state data assimilation (SDA) methodology for estimating process error covariance, which heretofore we refer to as process uncertainty. Our method uses sequential SDA, an iterative statistical approach that corrects process-model based predictions with field collected data at each time step and restarts the process-model with an update of the ecological variables of interest given from the data. Traditional sequential SDA approaches assume that the amount of process uncertainty contributing to total forecast uncertainty is known (Kalman, 1960), or that process uncertainty is proportional to observation error (Anderson et al., 2009); neither assumption is realistic for ecological systems. To address this limitation, we extend existing approaches to incorporate an estimate of the process uncertainty (i.e., the difference between the true state of the system and the forecast).

To make our approach more concrete, we consider the long history of forest gap modeling in ecology (Botkin et al., 1972; Solomon et al., 1980; Pacala et al., 1993; Post and Pastor, 1996), focusing on prediction of forest stand development at a single site and determining dominating uncertainty components. A forest gap model represents forest stand development arising from the birth, growth, and mortality of individual trees competing for light, water, and nutrients at the plot level, which is around 30 m^2^ (Bugmann, 2001). We first estimate model process uncertainty by assimilating 50 years of species-level aboveground biomass data using our novel SDA algorithm at Harvard Forest into LINKAGES, a well established forest gap model that was one of the first to “link” aboveground forest structure and composition to belowground biogeochemistry (Post and Pastor, 1996). As a result of our SDA process, we expect that we will improve prediction accuracy of aboveground biomass from LINKAGES. We also constrain an unobserved state variable, soil carbon, by leveraging the covariance between total soil carbon and aboveground biomass which the model provides. We expect that LINKAGES will accurately represent aboveground biomass processes, but that because it has been historically difficult to observe and understand the link between aboveground inputs and long-term soil carbon accumulation (Todd-Brown et al., 2013) that our aboveground-only constraint will not provide enough information to fully constrain belowground carbon pools. After applying the SDA algorithm to estimate process uncertainty, we then determine the most important sources of uncertainty by performing variance partitioning analysis across eight hindcasts of aboveground biomass from 1960 to 2010 (e.g., backtesting) each sequentially adding new components of overall uncertainty. While adding additional sources of uncertainty will lead to increased hindcast variance, we also expect that these hindcasts will be a much more accurate representation of our confidence in forecasting 50 years of aboveground biomass change than forecasts run in the typical spin-up initial conditions, static parameter, and known process uncertainty approaches.

This box provides background on each type of uncertainty in ecological process models.

- **Internal demographic stochasticity** - Demographic stochasticity refers to the variability in population growth arising from random sampling of birth and deaths.
- **Internal State (Initial conditions)** - Initial condition uncertainty is the uncertainty associated with the initial state of a system. For example, the number, type, and size of trees in a plot at the start of a model run.
- **External Forcing (Drivers / covariates)** - Driver and covariate uncertainty is typically the uncertainty around external environmental forcings like temperature and precipitation.
- **Parameter** - Parameter uncertainty arises because of our imperfect knowledge about the parameters in a model’s equations. Parameter uncertainties can be estimated by calibrating models to experimental or observational data (LeBauer et al., 2013; Fer et al., 2018), but are often fixed to single values in terrestrial ecosystem models.
- **Process** - Model process uncertainty is a measure of the ability of the model structure to predict the latent “true” state of the system after accounting for observation errors in the data. Without an estimate of process error covariance (multivariate) or process variance (univariate) it is difficult to determine model completeness. This would be analogous to predicting with a regression model without considering its RMSE.

## 2 Materials and Methods

Our methods are divided into four main steps. First, we provide background on the process model LINK-AGES and on the data from New England that we use to parameterize, validate, and assimilate. Second, we develop a novel sequential state data assimilation algorithm, the Tobit Wishart Ensemble Filter (TWEnF), which allows us to avoid a problematic assumption in the commonly used ensemble Kalman filter (EnKF): that the process error is known. Third, we ran eight model scenarios that additively include demographic stochasticity, parameters, external drivers, and process error. We estimated the last of our model uncertainty components, initial conditions uncertainty, by initializing each of the above scenarios with either ‘spin-up’ initial conditions or data-derived initial conditions. Spin-up initial conditions are created by running the model until an equilibrium state is reached; whereas, data derived initial conditions are created by running the model until an equilibrium state is reached than constraining the first time point with field collected data. Finally, we use the state variable outputs from these eight scenarios to calculate the contribution of each uncertainty component to total uncertainty through variance partitioning.

All of the model analyses took place within the Predictive Ecosystem Analyzer (PEcAn, pecanproject. org), an online framework for assimilating data into ecosystem models (Dietze et al., 2013). The specific modules we used within PEcAn, besides the basic workflow, were: allometry, sensitivity analysis, and sequential state data assimilation.

### 2.1 Ecosystem model

LINKAGES (Post and Pastor, 1996) is a forest gap model that links the dynamics of aboveground demographic processes with below ground biogeochemistry. At an annual time step, LINKAGES calculates the birth, growth, and mortality of individual stems as stochastic species-level functions of four environmental factors: soil moisture, growing degree days, available light, and available nitrogen. A decomposition subroutine governs the transitions of belowground carbon and nitrogen pools arriving as litter cohorts and driven by degree days, soil moisture, and soil nitrogen availability (Supplemental Figure 1). We chose LINKAGES as our process model because it efficiently captures the annual constraints on growth and mortality that match the tree-ring and census data we use to constrain the modeled stand dynamics of our study site, and because it has become an iconic depiction of forest gap processes (Bonan et al., 2019). While LINKAGES is our case study in this paper, our approach to data assimilation variance partitioning is generalizable to many process-based ecological models and data types.

A full analysis of the ecological dynamics inferred by LINKAGES at our site is beyond the scope of this paper and the subject of a separate manuscript (Raiho in prep.). In what follows, we focus on partitioning the total uncertainty in our estimates of aboveground woody biomass and soil carbon into the five uncertainties in Box 1. Aboveground woody biomass is a species-specific allometric function of stem diameter, which grows each year as a stochastic function of the most limiting of four environmental factors for each stem at an annual time step. In this study, modeled biomass increment is constrained by the assimilation of empirical estimates of aboveground woody biomass increment (See section 2.2). Soil carbon in LINKAGES is a stochastic function of the annual decomposition of litterfall cohorts that depends on respiration, which is itself a function of the ratio of lignin to nitrogen and actual evapotranspiration (Post and Pastor, 1996). We did not empirically constrain belowground state variables directly. Constraints on soil carbon, in this study, are indirect through the incoming source to litter pools, aboveground biomass.

### 2.2 Data sources and study site

We modeled the stand dynamics of the Lyford Plot, a 2.9 ha repeat-survey study site at Harvard Forest in central New England, USA (Foster et al., 2013) (42.53°N, 72.18°W). The stand initiated around 1900 following a prior history of grazing and logging. The stand lost chestnut in the 1910s due to the chestnut blight, was severely damaged by a hurricane in 1938, and experienced severe defoliation from a gypsy moth outbreak in 1981. The species that currently dominate the stand are mature red maple, which is typical of the region, and mature red oak, which is found in greater abundance in the stand than is regionally typical. The permanent plot was established by Walter Lyford in the 1960s and the diameter at breast height (DBH) and location of all stems over 5 cm DBH have been recorded at approximately decadal intervals since then. Additional site information including census collection is available in (Eisen and Plotkin, 2015).

Our model of stand development at the Lyforfd Plot includes the five species which currently make up 98% of the stems in the plot: red oak (*Quercus rubra*), red maple (*Acer rubrum*), yellow birch (*Betula alleghaniensis*), American beech (*Fagus Granifolia*), and eastern hemlock (*Tsuga canadensis*). There are 21 parameters per species in LINKAGES. To set Bayesian prior distributions for these parameters in our runs, we conducted a Bayesian meta-analysis to identify the parameters most needing constraint (LeBauer et al., 2013). We subsequently constrained the prior distributions of species-level specific leaf area (SLA) using trait data from the BETY database (LeBauer et al., 2018), and of allometric and recruitment parameters based on the literature (Catovsky and Bazzaz 2000; Dietze and Moorcroft 2011; Sullivan et al. 2017 in Supplemental part 2). These distributions can be found in the Supplemental Materials Section 2.

We validated the aboveground woody biomass produced by free runs of the model (before data assimilation) using biomass estimates from the Harvard Forest Environmental Measurement Station (EMS) Eddy Flux Tower (Munger, 2018) that is located approximately 2.4 km to the west of the Lyford Plot. The validation data are from DBH measurements collected annually since 1994. Allometric models were applied at the species-level using the Predictive Ecosystem Analyzer (PEcAn) allometry module (https://github.com/PecanProject/pecan/tree/master/modules/allometry).

Starting in the year 1960, we used empirical estimates of annual biomass increment data from the Lyford plot to constrain model runs via state data assimilation (see below). Our empirical estimates of biomass increment estimates and associated uncertainty derived from a Bayesian hierarchical model informed by annual growth increments from tree ring data and DBH at time of coring from the Lyford Plot (Dye et al., 2016), as well as from DBH values and tree status the decadal plot resurveys (Dawson et. al In Prep). To scale from estimated tree size to total aboveground biomass, taxon-specific allometric equations derived from Chojnacky et al. 2014 were used.

The meteorological drivers for our model runs were an ensemble (n=89) derived by probabilistically downscaling meteorological variables (temperature and precipitation) from global circulation models used in the Climate Model Intercomparison Project 5 (CMIP5) using a North American Land Data Assimilation System (NLDAS) training dataset (0.125 degree, hourly resolution), spanning from January 1979 to present (Xia et al., 2012). We used this met product instead of local meteorological data to generate a realistic representation of what driver uncertainty would be when making future predictions and to be consistent with other data assimilation runs being conducted by the Paleoecological Observatory Network (PalEON) (Rollinson et al., 2017). More information about these data sources and how they were processed can be found in the Supplemental Materials Section 1.1.

### 2.3 State data assimilation

We used sequential state data assimilation (SDA) to update species aboveground woody biomass at the end of each year from 1960 to 2010. While our analysis was a hindcast, we refer to it as a forecast because we ran the model in forward mode, only using data to validate our forecast, *post hoc*. Our process of sequential SDA followed the three steps of the forecast-analysis cycle (Dietze 2017a), repeated annually: 1) Forecast - An ensemble of LINKAGES runs (n=89) was used to make a probabilistic forecast of all the model’s state variables; 2) Analysis - We performed an SDA analysis of LINKAGES state variable prediction (described below). The state variables we assimilated were the biomass of five tree species and soil carbon amount; and 3) Update - We restarted LINKAGES with new state variable quantities leveraging the updated information (Supplemental Materials Section 5).

#### 2.3.1 Tobit Wishart Ensemble Filter (TWEnF)

We created a new SDA algorithm called the Tobit Wishart ensemble filter (TWEnF) to account for a non-normal likelihood and to estimate process uncertainty associated with LINKAGES during the analysis step. Our method is based on the ensemble Kalman filter (EnKF, Evensen 2009). The EnKF assumes that the forecast and observations both follow multivariate normal distributions, which allows it to have an analytical solution and operate efficiently. The normal assumption of the EnKF is often violated with ecological contexts, for instance, where a species might be locally absent while regionally present, or where a species may go extinct in a particular model ensemble while abundant on average across ensembles (Martin et al., 2005; Hall, 2000). We addressed this common problem by incorporating a Tobit likelihood into our analysis step (Figure 1, blue). Furthermore, while the analytical solution is computationally practical, the EnKF must make the assumption that process error is known. To estimate process error with data, our TWEnF introduces a latent ‘true’ state (Berliner, 1996) in the usual framework of Bayesian hierarchical models (Figure 1, pink).

**Figure 1:**
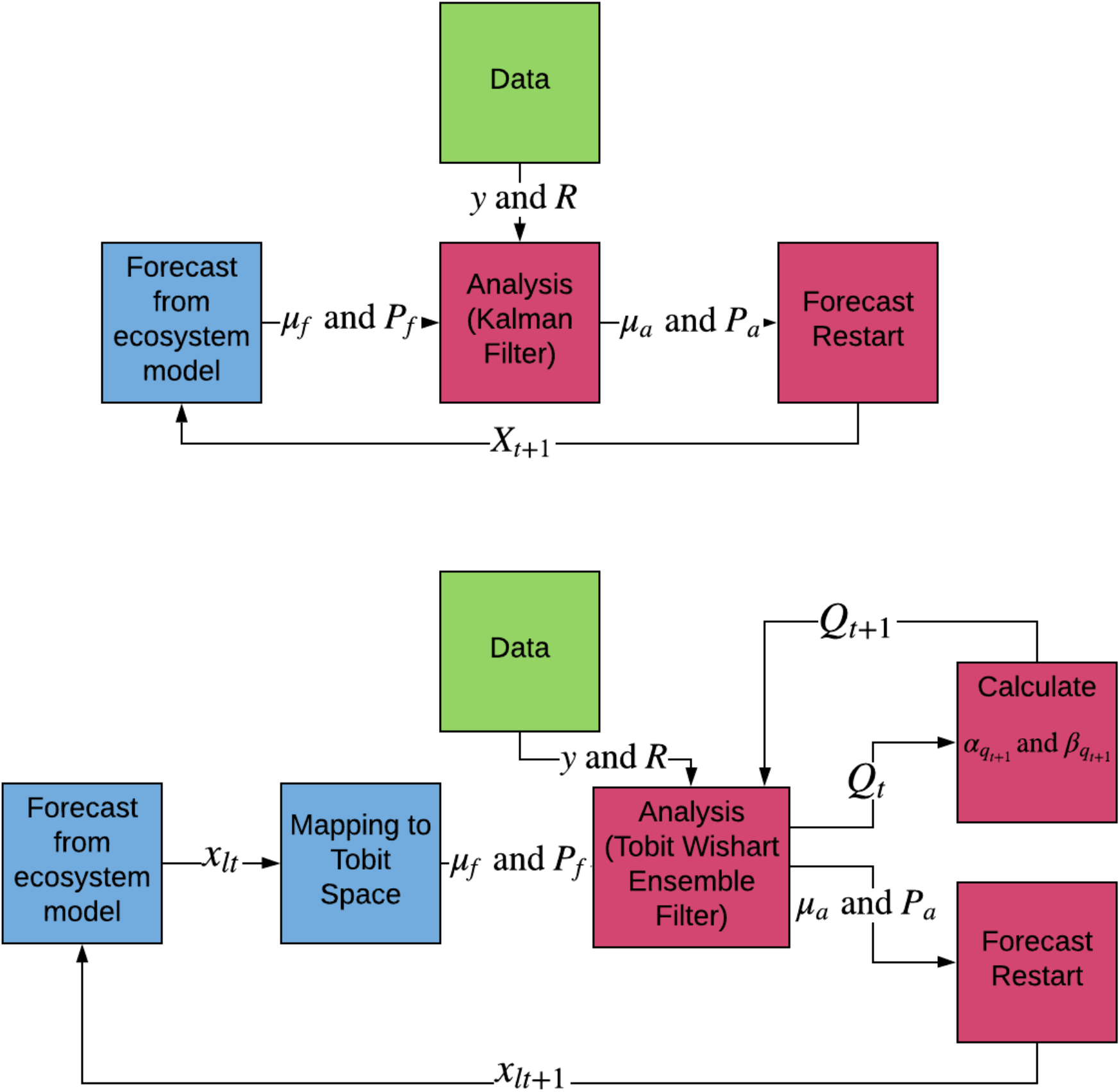
Conceptual diagram of the workflow involved in both the ensemble Kalman filter (EnKF, top) and the Tobit Wishart ensemble filter (TWEnF, bottom). Each method works in an iterative forecast cycle (Dietze 2017a) over time (*t* to *t* + 1), where the model forecast (blue) is updated by the data (green) into an analysis (pink), which is used to restart the forecast for another time step. The difference between these filters is that the TWEnF is generalized for non-normal forecasts and can also estimate the process covariance matrix over time by updating prior parameters (*α*_*q*_ and *β*_*q*_) at each time step. Let *y* be data mean, *R* be data covariance, *μ*_*f*_ be forecast mean, *P*_*f*_ be forecast covariance, *μ*_*a*_ be mean analysis, *P*_*a*_ be analysis covariance, *Q*_*t*_ be process covariance matrix at time *t*, and *x*_*lt*_ be the left censored ecosystem model ensemble values. In both cases, the analysis analysis mean (***μ***_*a*_) and covariance (***P***_*a*_) are taken from the filter and used to update the ecosystem model states which restart the next ecosystem model forecast.

To estimate process error covariance, we fit the following TWEnF annually to tree-ring derived multivariate species biomass (***y***) informed by prior information from the calculated mean (***μ***_*f*_) and covariance (***P***_*f*_) of the model ensemble (n=89). We modified the likelihood to account for zero-truncated (or left-censored) data (eqn. 1) then we estimated process error covariance (***Q***), where

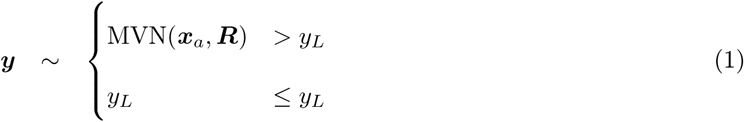

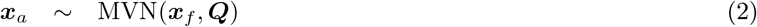

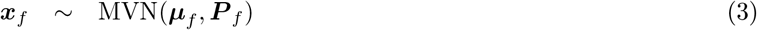

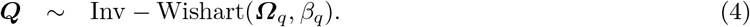

Let ***y*** be the posterior mean of multivariate species biomass from the aforementioned tree ring analysis; ***R*** be, similarly, the posterior covariance of species biomass from the tree ring analysis; *y*_*L*_ be the left censored threshold which is equal to 0 in our case; ***x***_*a*_ be representative of the true multivariate species biomass state; (***Q***) be the covariance of the latent state (***x***_*a*_), with mean (***x***_*f*_) arising from the forecast ensemble mean (***μ***_*f*_) and covariance (***P***_*f*_). We mapped ***μ***_*f*_ and ***P***_*f*_ to Tobit space in a previous step that incorporates known meteorological weights (Papadakis et al., 2010) using a very similar model formulation to the TWEnF, which is described further in Supplemental Materials Section 4. Finally, we calculated the analysis mean (***μ***_*a*_) and covariance (***P***_*a*_) used to restart the ecosystem model as a derived quantity from the estimated latent state (***x***_*a*_).

We updated the estimate of the process covariance (***Q***_*t*_) every time step by updating the shape parameters of the Inverse Wishart distribution as follows

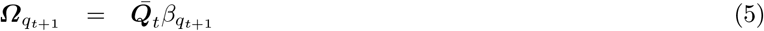

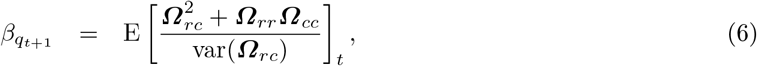

where ***Ω*** is the process precision to the process covariance ***Q***_*t*_ and *r* and *c* represent the rows and columns of the process precision matrix. In this step of the analysis, 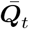 is the posterior mean estimate of ***Q***_*t*_ from the TWEnF. We assessed convergence at every time step using the Gelman-Rubin convergence diagnostic in the ‘coda’ package (Gelman et al. 1992; Plummer et al. 2006) over three MCMC chains of 100,000 iterations each. We restarted our workflow with different initial conditions if Gelman-Rubin diagnostics were greater than 1.01 for more than two monitored variables.

### 2.4 Hindcasting uncertainty scenarios

We used eight model scenarios with additively more types of uncertainty to partition total forecast variance between the five components we considered (Box 1). In order to partition uncertainty from initial conditions, we divided the scenarios into two initial condition types: model spin-up (run without data constraints until equilibrium is reached, Scenarios A1-4) or informed by data (after spin-up the forecast is constrained with data from tree ring derived biomass in 1960 Scenarios B1-4). In each scenario batch (A versus B), initial condition uncertainty was the first type of uncertainty added to the default ecosystem model. LINKAGES includes demographic stochasticity by default, so the default version of LINKAGES plus initial condition uncertainty were scenarios A1 and B1 in this analysis. We then added the following uncertainties sequentially: parameter (A2 and B2), meteorological (A3 and B3), and process uncertainty (A4 and B4) (Table 1). Parameter and meteorological uncertainties were added by running each ensemble member with a different parameter and meteorological set, sampled from the calibration posteriors and meteorological ensemble. By contrast, in the default runs (A1 and B1), all ensemble members were run at the posterior means. Finally, to incorporate process uncertainty we used the final posterior mean of the process error covariance (*Q*) from the full data assimilation run described in Section 2.3. In this scenario, runs were conducted leveraging the forecast-analysis cycle, stopping the model each year to add process error then restarting the process-model, but no data constraints were added during the analysis step (except for year 1 in the B1, data constrained initial conditions scenario).

**Table 1:** The types of uncertainty included in each scenario are indicated with an ‘X.’

| Scenario | Demographic | Initial Conditions | Parameter | Meteorological | Process |
| --- | --- | --- | --- | --- | --- |
| A1 | X | Spin Up |  |  |  |
| A2 | X | Spin Up | X |  |  |
| A3 | X | Spin Up | X | X |  |
| A4 | X | Spin Up | X | X | X |
| B1 | X | Data Derived |  |  |  |
| B2 | X | Data Derived | X |  |  |
| B3 | X | Data Derived | X | X |  |
| B4 | X | Data Derived | X | X | X |

### 2.5 Variance partitioning

Variance partitioning allows us to quantify which aspects of uncertainty contribute the least to overall uncertainty and pinpoints where we should focus efforts to constrain uncertainty in future predictions. We estimated the effect of each source of variance by calculating the difference in variance between pairs of scenarios then calculating the cumulative proportion of variance in reference to the final scenario that includes all five aspects of uncertainty (Dietze, 2017b). This is similar to analytical approximation methods (Hawkins and Sutton, 2009) but our sequential approach accounts for nonlinear interactions that may affect prediction.

Our scenarios did not allow a full variance partitioning because we did not introduce each source of variance independently from the other sources of variance in all possible permutations. Specifically, it was not possible to partition initial condition variance because we could not separate initial condition uncertainty from demographic stochasticity. However, because we had two sets of scenarios: one with data derived initial conditions (scenarios A1-4) and one with spin-up based initial conditions (scenarios B1-4), we were able to calculate the covariance between data derived initial condition variance and the other components of variance (Cov[*A, B*], eqn. 7). This calculation shows the duration and magnitude of the impact of data derived initial conditions on the forecast variance. As an example, we calculated the magnitude of the interaction terms with the following equation for variance between two variables P (parameters) and IC (initial conditions):

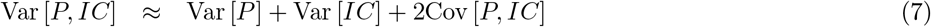

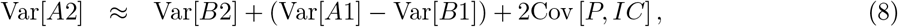

where we substitute Var[*P, IC*] with the variance from scenario A2 (Var[*A*2]). Scenario A2 includes uncertainty from spin-up and parameters. We then also substitute Var[*P*] with variance from scenario B2 (Var[*A*2]). Scenario B2 includes uncertainty from parameters and constrained initial conditons. Finally, we also substitute Var[*IC*] with the difference between variance in scenarios A1 (spin-up) and B1 (constrained IC) where neither include uncertainty from parameters. We used these values to solve for Cov[*P, IC*], which is the covariance between initial condition uncertainty and parameter uncertainty. Following similar logic, we can solve for Cov[*M, IC*] and Cov[*Process, IC*] using the difference between the subsequent scenario variances as the Var[*IC*].

## 3 Results

### 3.1 Model Parameterization

We found that running LINKAGES using the default parameters resulted in inaccurate predictions of forest composition and biomass when compared with species-level biomass data from the nearby Harvard Forest EMS Tower (Munger, 2018). Free runs of LINKAGES using data constrained parameters improved the accuracy of predicted total biomass but not that of forest composition (Figure 2 and Table 2). Under default parameterization, LINKAGES predicted that hemlock would be the dominant species, and the stand was predicted to have low total stand biomass (*≈* 5 kgC/m^2^). As is the case at the Lyford plot, red oak was the dominant species at the EMS tower plot, and the site had higher total stand biomass (*≈* 15 kgC/m^2^). Parameters informed from our specific leaf area (SLA) meta-analysis (LeBauer et al., 2013) and literature review for allometric and recruitment parameters (Catovsky and Bazzaz, 2000; Dietze et al., 2008; Sullivan et al., 2017) allowed LINKAGES to better represent total forest biomass (*≈* 10kgC/m^2^) but not species composition (Euclidean Distance = 11.52 versus 10.89 with default parameterization). After informing parameters with independent data, our parameter uncertainty analysis revealed that there were some parameters that could be constrained with model calibration, but that the majority were not causing sufficient model sensitivity to warrant a full parameter calibration effort (See supplemental section 3.1).

**Figure 2:**
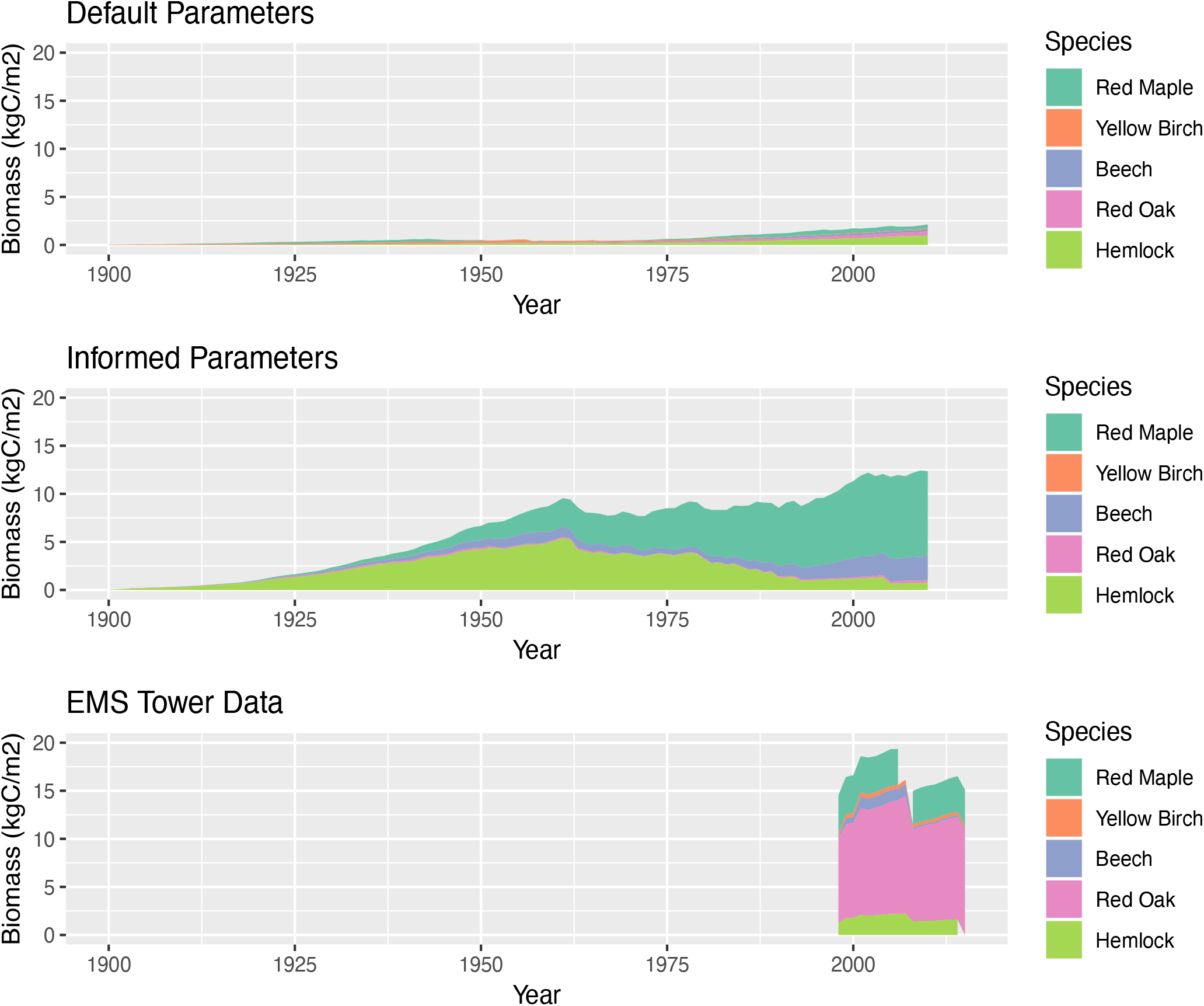
Cumulative time series of species-level biomass from LINKAGES run with default parameters (panel 1) and parameters derived from informative priors (panel 2). We compare these results with tree diameter at breast height (DBH) data collected from the trees surrounding the Harvard Forest EMS tower (Munger 2018, panel 3).

**Table 2:**
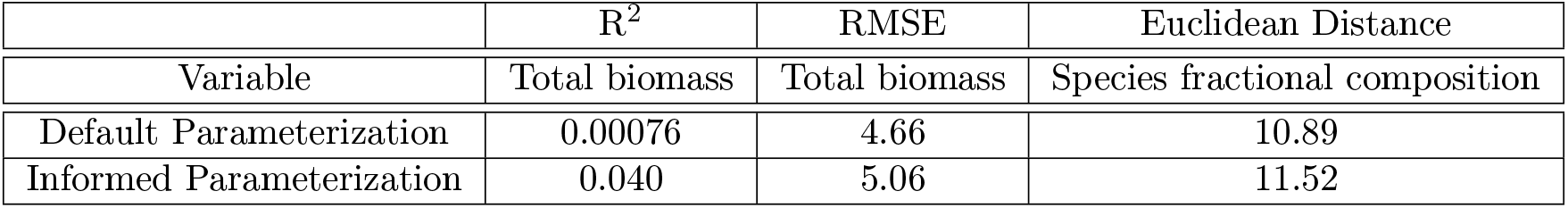
Bias diagnostics for the output of LINKAGES free runs compared with Harvard Forest EMS Tower data showing results from the default and calibrated parameterizations. R^2^ and root mean square error (RMSE) are calculated for total stand aboveground woody biomass and Euclidean distances (e.g., root square sums) are calculated between average species composition vectors over the time period 1998 to 2009.

| | $R^2$ | RMSE | Euclidean Distance |
| --- | --- | --- | --- |
| Variable | Total biomass | Total biomass | Species fractional composition |
| Default Parameterization | 0.00076 | 4.66 | 10.89 |
| Informed Parameterization | 0.040 | 5.06 | 11.52 |

### 3.2 State data assimilation

Empirical estimates of aboveground biomass derived from tree ring and census observations at the Lyford plot showed that red oak, the dominant species in the stand, has accrued biomass over the last fifty years while understory species have experienced a few mortality events among individuals. The census was not conducted annually, therefore the biomass data has larger uncertainty during periods where an individual in the understory has died. For example, the green envelope spanning the data in Figure 3 had high uncertainty between 1980 and 1990, a census period that experienced both yellow birch and hemlock mortality. These areas of larger uncertainty allowed us to illustrate an example of successful constraint by our methods, as the analysis step (Figure 3, pink) was able to match the variance associated with the data during those time periods. We assessed our state data assimilation algorithm by looking at several bias diagnostics: average model bias (difference between the observation and analysis over time), mean square error, R^2^, relative absolute error, and absolute mean error (Supplemental Section 6). We also reported the coefficient of variation for the average model bias to account for differences in species biomass magnitudes. The highest biomass species, red oak, was best represented by LINKAGES with a high R^2^ between the modeled red oak and the data (R^2^= 0.769) (Supplemental Figure 5). As the most abundant species, red oak unsurprisingly had the largest average model bias (−0.82 kgC/m^2^/yr, 10.86% coefficient of variation (CV), Table 3) and largest estimated process variance (diagonal element of process error covariance matrix) among the aboveground biomass of species (*σ*^2^ = 0.25, 14.2% CV). This bias increased over time, indicating that the modeled process of red oak mortality and/or growth may need adjustment and agreeing with ecological analyses that red oak will continue growing at Harvard Forest in the future (Eisen and Plotkin, 2015).

**Figure 3:**
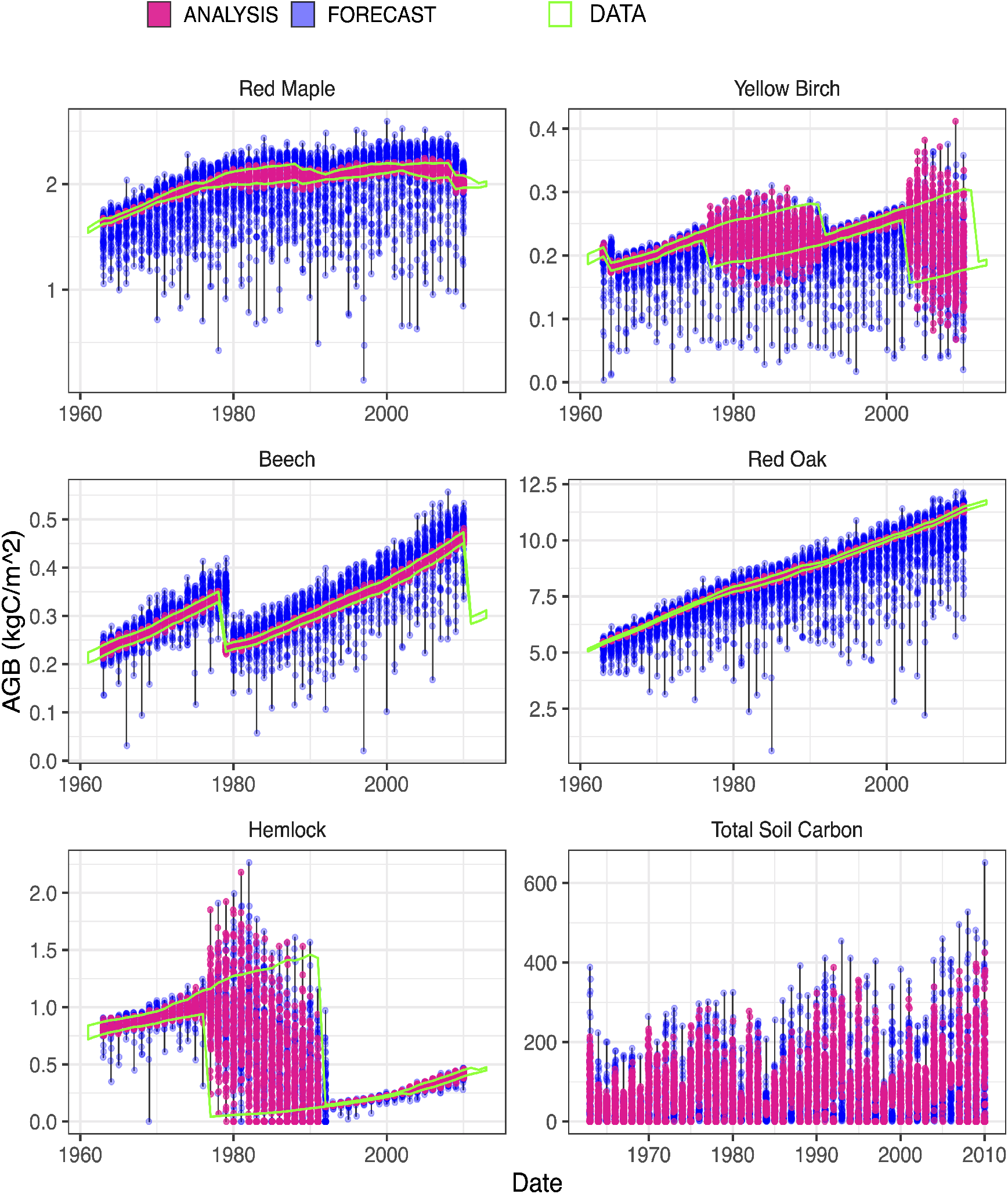
Species biomass time series illustrating the difference between the model forecast ensembles (blue points), the data 95% credible intervals (green lines), and the analysis ensembles (pink points) in LINKAGES state data assimilation of tree ring derived aboveground biomass. The green confidence intervals in front of the pink and blue points are credible intervals of the tree ring estimated species level biomass. The black vertical lines indicate time points where data was assimilated: annually between 1961 and 2010. The blue points are 89 LINKAGES forecasts of one year forward following an analysis (pink). The pink points are 89 species biomass values drawn from the estimates of average species biomass in the Tobit Wishart ensemble filter (TWEnF). The analysis points are used to restart the 89 model ensemble members for the next cycle of annual forecasting. The pink points generally align with the mean of the data while the forecasts sometimes drift from the data. During the time span between some censuses, the data are bimodal and appear to show wide uncertainty because the timing of mortality events within these census intervals is unknown.

**Table 3:** Model diagnostics ordered by species biomass. Average model bias is simply the modeled mean minus observed annual means. Estimated process variance (diagonal elements of process error covariance matrix) for each species is estimated over time using the Tobit Wishart ensemble filter (TWEnF). Average forecast total biomass is the average modeled biomass for each state variable to give a reference point for the magnitude of the estimated process variance. For example, red oak is the highest biomass species in the stand and also has the highest estimated process variance.

| State Variable | Average Model Bias (kgC/m<sup>2</sup>/yr) | Estimated Process Variance | Average Forecast Total Biomass (kgC/m<sup>2</sup>/yr) |
| --- | --- | --- | --- |
| Red Oak | -0.82 | 0.25 | 7.55 |
| Red Maple | -0.14 | 0.13 | 1.86 |
| Eastern Hemlock | 0.06 | 0.13 | 0.62 |
| American Beech | -0.04 | 0.13 | 0.28 |
| Yellow Birch | -0.01 | 0.09 | 0.22 |
| Total Soil Carbon | – | 1.03 | 9.83 |

The second most abundant species, red maple, had a persistent negative bias (−0.14 kgC/m^2^/yr, Figure 2). This negative bias was expected because red maple is the dominant species in the region but is suppressed at Harvard Forest by regionally anomalously large red oak (Lorimer, 1984; Abrams, 1998). Both red oak and red maple’s negative biases were consistent with the unconstrained (‘free’) run (where oak and maple went locally extinct) and suggested a need for red oak and red maple parameter calibration and/or evaluation of the ecological competitive process in LINKAGES (Figure 2). However, both species had low estimated process variances (*σ*^2^ = 0.25, 0.13 and 14%, 7% of total process uncertainty respectively, Table 3) indicating that LINKAGES modeled representations of the two most abundant species were adequate for prediction so long as parameter uncertainty and process covariances are incorporated in the analysis.

We estimated the process covariance matrix, akin to RMSE in linear models (Box 1), associated with a process-based ecological model. Linearly increasing posterior estimates for the process covariance matrix degrees of freedom over time (Figure 4 left) provided evidence that the estimation of the process covariance matrix was increasingly constrained over time and could continue to be constrained by a longer time series of data. The values associated with the biomass of each species, along the diagonal of the process covariance matrix, were estimated to be small (Table 3 column 3), indicating that annual species biomass accumulation process was well represented by the forest gap model (Shugart et al., 2020), once we accounted for uncertainty in the data. Similarly, the species correlations in the process covariance matrix were estimated to be small with the most significant correlation between species being a small negative relationship between beech and red oak (correlation = −0.125, Figure 4). This suggested that, while LINKAGES typically represents beech and red oak as having a positive interaction (forecast correlation = 0.105, Supplemental Materials Figure 9), they were actually more neutral with one another at Harvard Forest according to the tree ring data.

**Figure 4:**
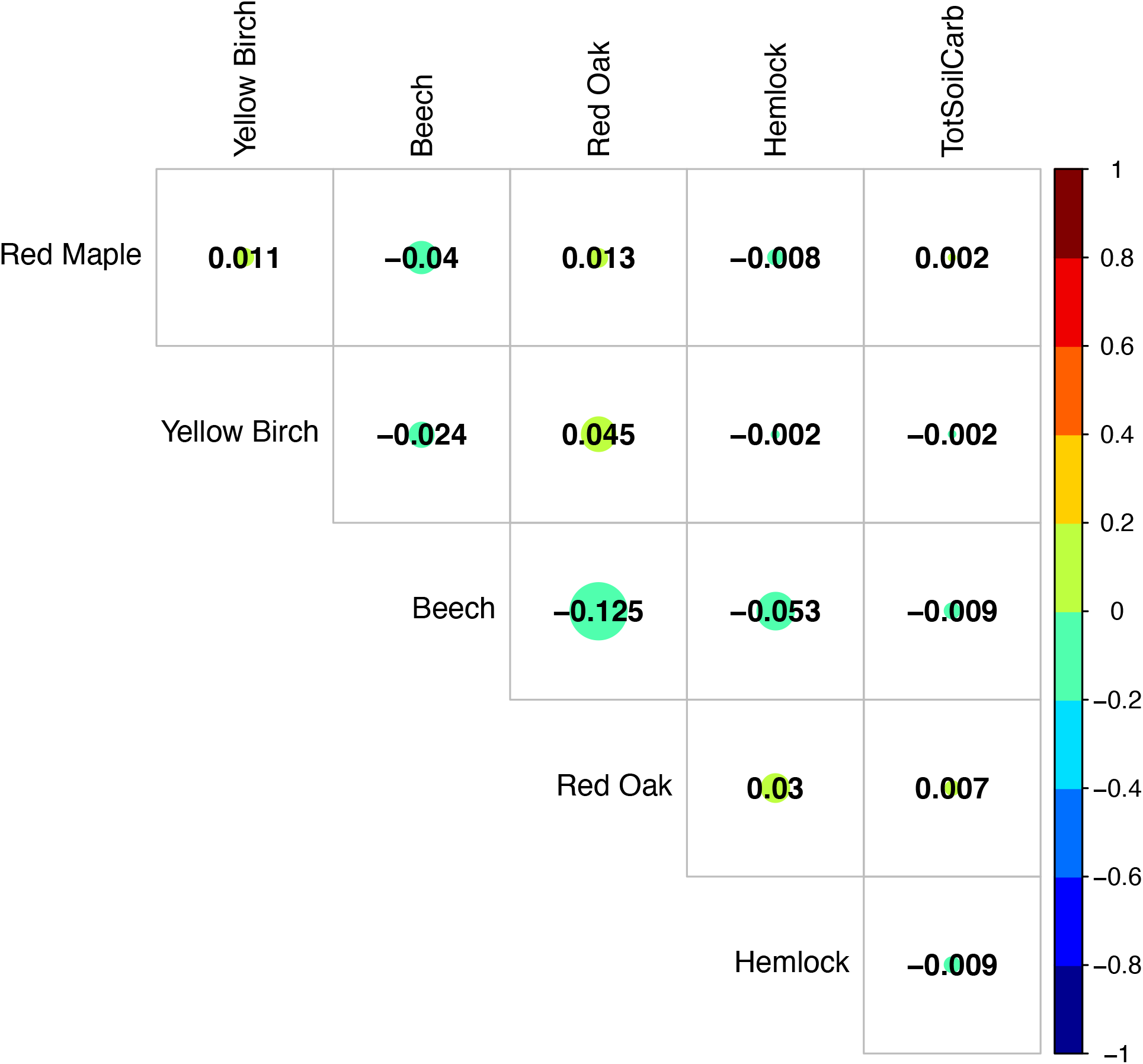
A correlation diagram of the process covariance. The colors in the correlation diagram correspond to the magnitude and direction of the correlation. The diagonal variances can be found in Table 3.

In the absence of empirical data on changing soil carbon pools, we depended on the mechanistic linkages between aboveground and belowground carbon in LINKAGES to constrain soil carbon fluxes. In our runs, LINKAGES did not provide a constraint on soil carbon given the aboveground biomass constraint and soil carbon pools rose to highly unrealistic levels with a similarly high process variance estimate (*σ*^2^ = 1.03, Table 3). Soil carbon was not estimated to be highly correlated with any species biomass in the process error covariance matrix (correlations between −0.009 and .0002), but was somewhat more correlated with species biomasses in the forecast ensemble covariance matrix (correlations between −0.238 and 0.168, Supplemental Figure 9). The lack of constraint on the flux of soil by aboveground dynamics in our results is puzzling, and may reflect undetected errors in our version of the process model.

### 3.3 Hindcasting uncertainty scenarios

Across all uncertainty scenarios, data constrained initial conditions reduced model bias and improved root mean square error agreeing with ecological hypotheses that forests and potentially many ecological processes have substantial historical dependence that should be accounted for by propagating initial condition uncer-tainty. Across most uncertainty scenarios, excluding scenario A1 and B1, data constrained initial conditions lowered average forecast variance (Table 4). Scenarios A1 and B1, the default model run with only demographic stochasticity and initial condition uncertainties alone (Figure 5, row 1), were precise (average forecast standard deviation = 0.32 kgC/m^2^ and 1.450 kgC/m^2^) compared to the actual residual error (average observation standard deviation = 0.27 kgC/m^2^). However, the root mean square error between the hindcast and the data decreased when the initial conditions were constrained with data (RMSE = 9.81 spin-up initial conditions, 4.42 data derived initial conditions; Table 4). The correlation coefficient was closer to one in scenario A1 (0.934) versus scenario B1 (0.262) because without data constrained initial conditions biomass increases more linearly.

**Table 4:** Model diagnostics for hindcasting scenarios. Data constrained initial conditions reduce model bias across scenarios. For average model bias, a value closer to zero indicates less bias. For the correlation coefficient, a value closer to 1 indicates stronger correlation between the predictions and the data. For root mean square error, a lower value indicates a smaller difference between predictions and data. Average forecast variance increases as we add more types of uncertainties as expected.

| Hindcast Diagnostic | Scenario | Demographic Stochasticity | + Parameter | + Meteorological | + Process |
| --- | --- | --- | --- | --- | --- |
| Average Model Bias (kgC/m<sup>2</sup>/yr) | A1: Spin-up IC | -9.72 | -4.85 | -2.89 | 4.090 |
| Average Model Bias (kgC/m<sup>2</sup>/yr) | B1: Data IC | -4.07 | -2.23 | -1.88 | 0.669 |
| Correlation Coefficient | A2: Spin-up IC | 0.934 | 0.972 | 0.968 | 0.966 |
| Correlation Coefficient | B2: Data IC | 0.262 | 0.912 | 0.929 | 0.954 |
| Root Mean Square Error | A3: Spin-up IC | 9.81 | 4.86 | 3.29 | 5.39 |
| Root Mean Square Error | B3: Data IC | 4.42 | 2.35 | 2.01 | 2.00 |
| Average Forecast Variance (kgC/m<sup>2</sup>/yr) | A4: Spin-up IC | 0.103 | 141.0 | 310.0 | 326 |
| Average Forecast Variance (kgC/m<sup>2</sup>/yr) | B4: Data IC | 1.450 | 20.3 | 30.1 | 71 |

**Figure 5:**
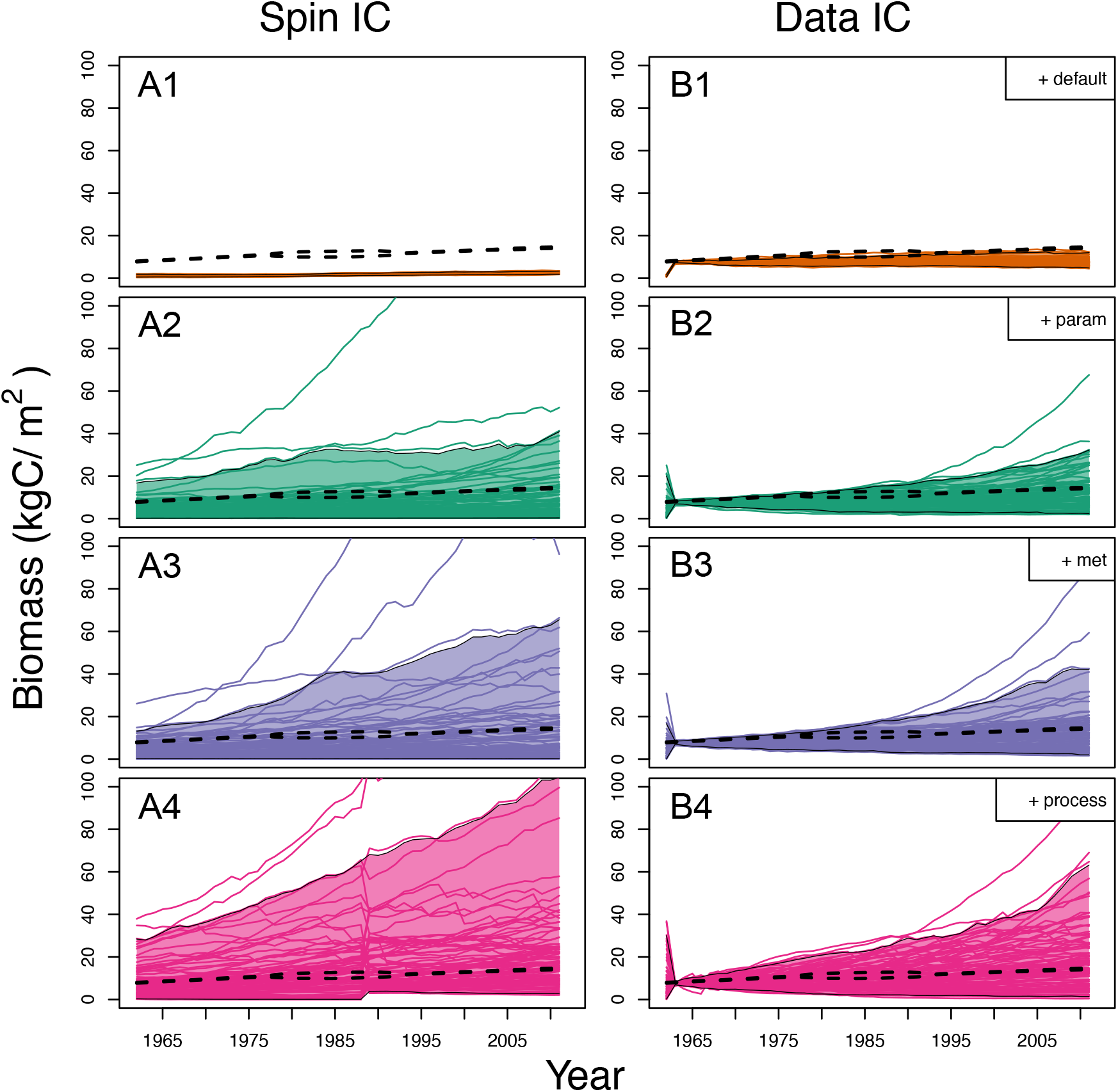
Individual model ensemble members overlaid with shaded 95% quantiles (outlined in black) of aboveground biomass results from each uncertainty scenario using spin-up as the initial conditions in the model (left) and using data to constrain the initial conditions in the model (right). Default was run with initial condition uncertainty and internal model demographic stochasticity, which vastly under-represents the true forecast uncertainty (first row). Next, parameter uncertainty was accounted for (second row), followed by meteorological uncertainty (third row). Finally process uncertainty estimated in the full data assimilation was accounted for (fourth row). The dotted lines on all the plots are the 95% credible intervals of the data estimated from tree rings.

To represent a full characterization of the state of knowledge of the system, we sequentially accounted for uncertainties (scenarios A2-4 and B2-4, Table 4), which illustrated that the true variance in our forecast is large. The variance in the spin-up scenarios (A1-4) was consistently larger than the data constrained initial conditions (B1-4, Table 4). Accounting for meteorological uncertainty increased variance in the spin-up initial conditions (variance increased *≈* 170 kgC/m^2^/yr) much more than the data-derived initial condition scenario (variance increased *≈* 10 kgC/m^2^/yr) (Figure 5, row 3, Table 4). Recognizing process uncertainty allowed us to see see the substantial uncertainty in the spin-up initial condition scenario (Figure 5, column 1, row 4) and even more so in the data constrained initial condition scenario (Figure 5, column 2, row 4). Accounting for process uncertainty reduced model bias (from −1.88 to 0.669) and increased the correlation coefficient slightly between the model and the data (from 0.929 to 0.954) in the data derived initial condition scenarios. This improvement occurred because the process covariance constrained species biomass by inducing species covariances that were not present in the model but were present in the data (Figure 6, Supplemental Figure 6). In these scenarios, more model ensembles included sub-canopy red maple, yellow birch, and hemlock, which were less abundant in scenarios that do not include process covariance (scenarios A1-3 and B1-3).

**Figure 6:**
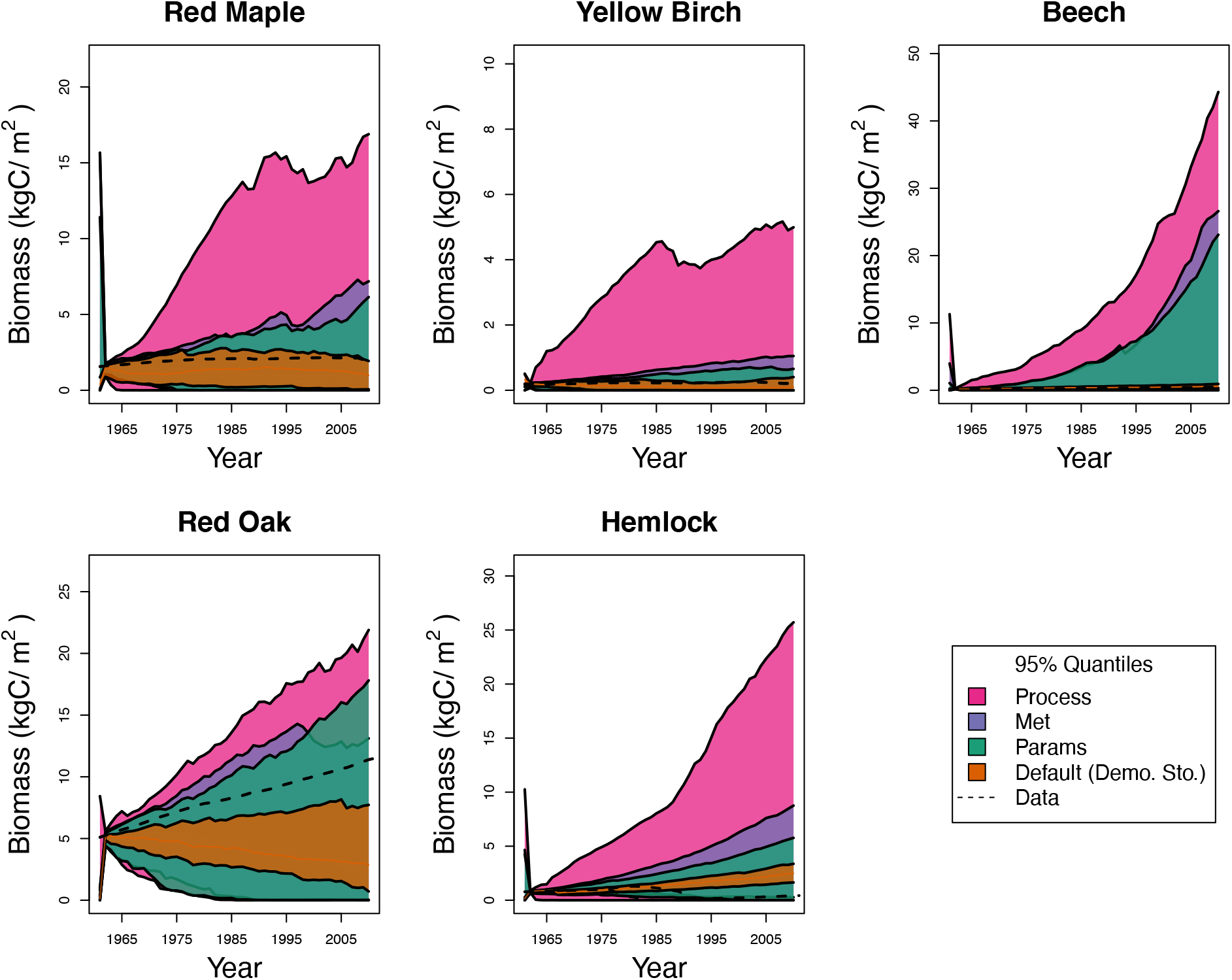
95% quantiles of species level biomass over time colored by the four data constrained initial condition uncertainty scenarios (scenarios B1-4). Uncertainty from ecosystem model spin up can be seen up to 1960 then data constraints greatly constrain the forecasts. Variance from the first scenario arises only from demographic stochasticity (orange). Parameter, meteorological, and process uncertainty are sequentially accounted for in the next three scenarios. The dotted lines on all the plots are species posterior means of the data estimated from tree rings. Note that the y-axes are different between plots to provide better visualization of the uncertainty components for lower biomass species.

### 3.4 Variance Partitioning

Variance partitioning showed that the covariance between the initial condition uncertainty and the other types of uncertainties was the dominant variance contributor over time (hashed areas in Figure 7). All model scenarios that were run with model spin-up had much larger uncertainty than with data constrained initial conditions (Figure 5, column 1 versus column 2). In addition, initial conditions had long lasting effects on the magnitude of the total forecast variance (Figure 7). Notably, covariance between initial condition uncertainty and parameter, meteorological driver, and process uncertainty decreased significantly over time while the interaction between initial conditions and demographic stochasticity slightly increased because inducing different stand types initially increased the variance in stand trajectory over time. Overall comparing scenarios A1-4 to scenarios B1-4 shows that a one time constraint on initial conditions was able to limit the total variance for 50+ years. The exponential decay constant of the effect of the initial conditions on total biomass variance was .08/year, meaning that the half life of the effects of initial condition constraint is 4.25 years. However, after 50 years the total forecast variance of the spin-up initial conditions was still 12.24% higher than the total forecast variance of the data constrained initial conditions.

**Figure 7:**
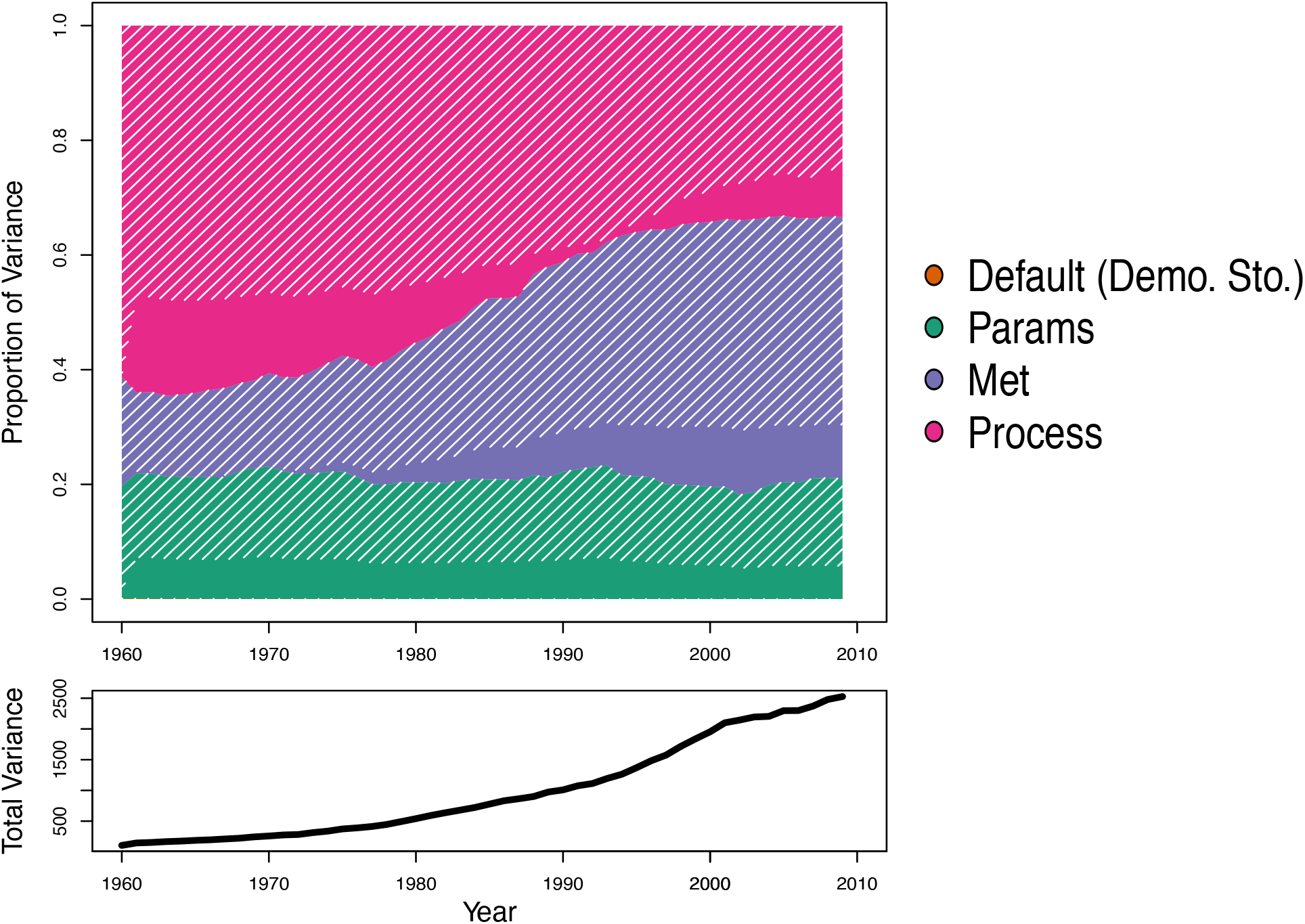
Top: The relative contribution of each type of variance to total aboveground biomass variance. The hashed areas are the relative variances that can be attributed to the covariance with initial conditions. For example, over time initial condition uncertainty covariance with meteorological uncertainty (purple) accounted for a larger proportion of total variance. Bottom: The black increasing line indicates the total amount of aboveground biomass (kgC/m^2^) variance partitioned by the relative variance plot. This shows that while the proportion of variance that process variance is contributing to the total variance decreases over time that the absolute magnitude of that variance is not necessarily decreasing.

Because we did not assimilate soil carbon data but updated soil carbon based mechanisms in the model (litterfall, mortality, decomposition, etc), we considered the soil carbon uncertainty separately from above-ground biomass uncertainty. The initial condition constraint was much less apparent in the soil carbon variance partitioning results outside of major outliers (Figure 8, left). Process uncertainty dominated by an order of magnitude, reflecting the lack of constraint by our version of LINKAGES on this carbon pool (Figure 9), which was out of the bounds of any soil carbon pool on Earth. Even though process uncertainty was the obvious contributor to total uncertainty, meteorological, and parameter uncertainty also caused total soil carbon to drift to extremely large values. The covariance between initial conditions and process uncertainty was an increasingly substantial component over time (Figure 8, right), but it was difficult to assess how much of a constraint initial conditions could provide given the magnitude of uncertainty for total soil carbon.

**Figure 8:**
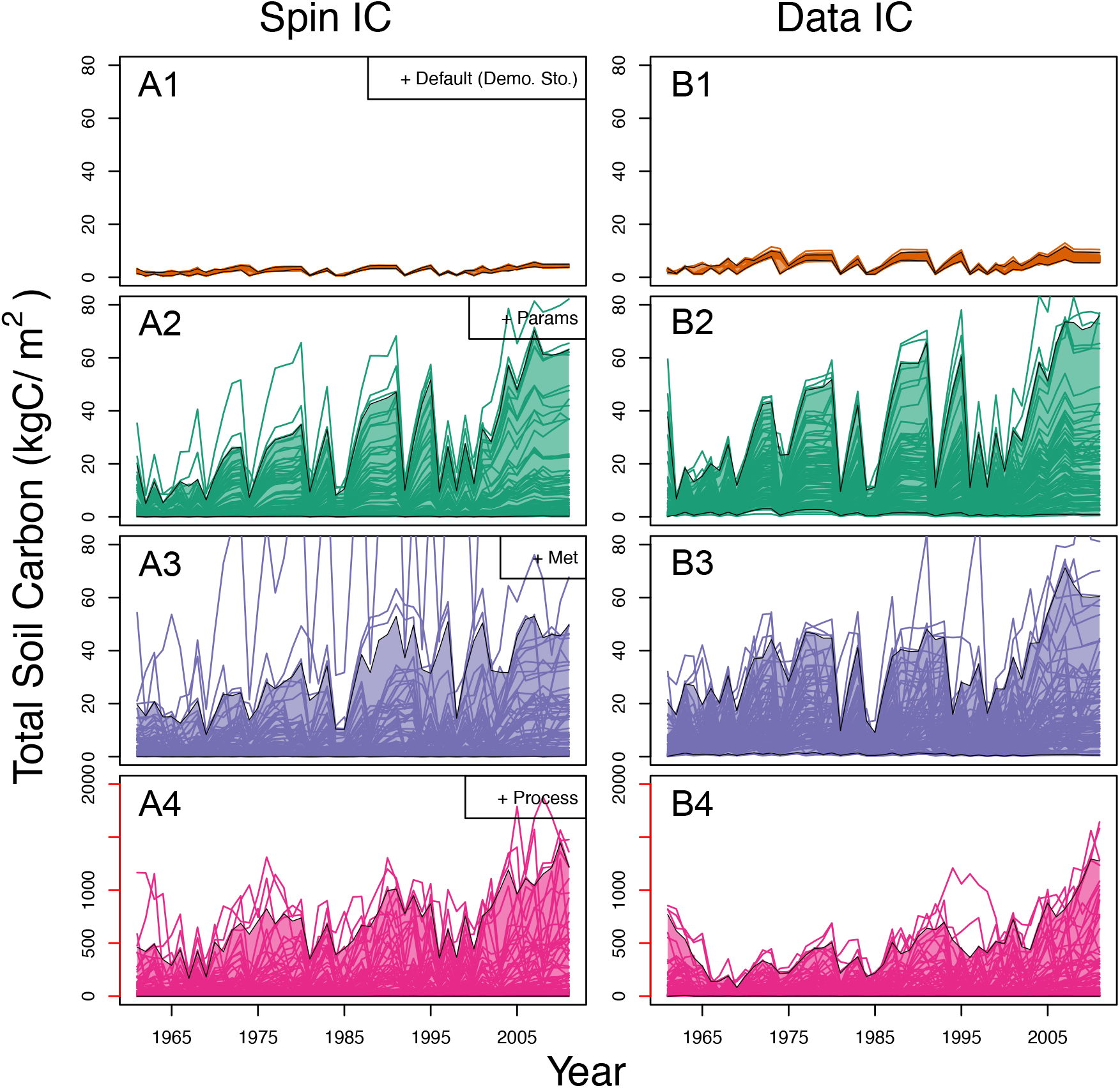
Individual model ensemble members overlaid with shaded 95% quantiles (outlined in black) of total soil carbon results from each uncertainty scenario using spin-up as the model’s initial conditions (left) and using data to constrain the model’s initial conditions (right). The default was run with initial condition uncertainty and internal model demographic stochasticity (first row). Next, parameter uncertainty was included (second row), followed by meteorological uncertainty (third row), and finally process error (fourth row). The y-axis in the fourth row is colored differently to draw attention to the much larger scale in this row. The instability in the soil carbon reconstruction arises from deterministic cohort dynamics present in the version of LINKAGES we ran.

**Figure 9:**
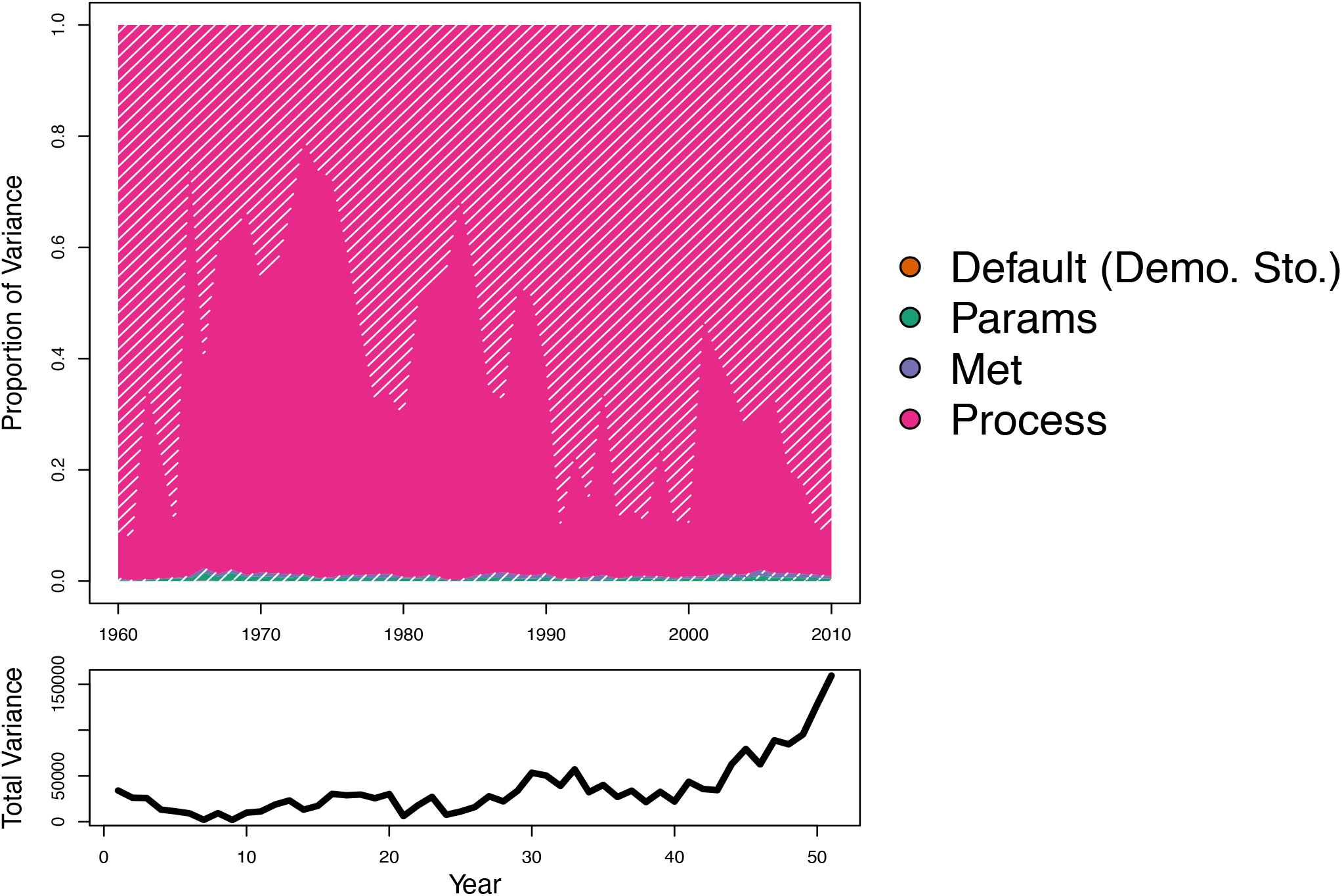
Top: The relative contribution of each type of variance to total soil carbon variance. The hashed areas are the amounts of variance that can be attributed to the covariance with initial conditions. Process (pink) uncertainties contributed a large amount of proportional variance to covariance with initial condition uncertainty. Bottom: The black increasing line indicates the total amount of variance in soil carbon (kgC/m^2^) that is being partitioned by the relative variance plot above.

## 4 Discussion

In our final modeling scenario (B4), we incorporated five data constrained uncertainties: demographic stochasticity, parameter, meteorological, initial condition, and process uncertainty (Figure 5 and 8, bottom right). This suite of uncertainties represents the current state of knowledge of a 50 year prediction of forest stand development at Harvard Forest provided by LINKAGES. While our quantification of the above-ground biomass trajectory of the Lyford Plot at Harvard forest is uncertain, this is an accurate depiction of the ability of LINKAGES to predict the biomass trajectory of a single stand. The five uncertainties discussed here are present, whether we estimate them or not.

Most predictions of forest succession, however, fail to quantify most of the uncertainties associated with their forecasts. We identified 15 papers published between 2008 and 2018 that used forest gap models explicitly for forecasting (Fischer et al. 2015, 2014, 2016; Gutiérrez et al. 2016; Morin et al. 2014; Sun et al. 2018; Taylor et al. 2017; Boulanger et al. 2018; Foster et al. 2017; Chauvet et al. 2017; Rödig et al. 2017b,a, 2018). Demographic stochasticity was included in all of them, but only Gutiérrez et al. (2016) accounted for any other type of uncertainty. Our study highlights the consequences of ignoring these uncertainties. As an illustrative example, in the top left panel of Figure 5, the default parameters of LINKAGES make precise predictions of the stand’s above ground woody biomass, which are well outside of the distribution of empirically observed AGWB. To address such a poor hindcast, modelers typically would ‘tune’ parameters until the model hindcast had improved and accept the tuned model as a reasonable estimate of the observed state variable (Bugmann et al., 2001). This approach misleadingly attributes model bias entirely to parameter uncertainty, distorts the prediction through *post hoc* analysis, and presents a forecast that is artificially precise and accurate (Wramneby et al., 2008). By systematically quantifying and partitioning total forecast uncertainty, we demonstrate that parameter uncertainty makes a relatively small contribution to the forecast of AGWB at Harvard Forest using LINKAGES. Instead, we found that the vast majority of uncertainty in the forecast is due to initial conditions and process uncertainty, two sources of variance that are rarely estimated or included in ecosystem modeling. A focus on the reduction of process and initial conditions uncertainty would represent a substantially different approach to improving the predictive power of gap models than the direction of the bulk of current research efforts and default modeling assumptions.

### 4.1 Demographic stochasticity, parameter, and meteorological uncertainty

Our results indicate that relying on demographic stochasticity alone to characterize the variability in ecological processes may result in overconfident and inaccurate model forecasts. Predictions using stochastic forecast gap models could be misleading scientists, managers, and policy makers because the model projections appear precise, while also appearing to account for uncertainty, but are only accounting for a tiny fraction (0.09%) of uncertainty in the projection in our case study.

Overemphasis on local parameter tuning may lead to overconfidence in predictions and will decrease the ability of a model to be generalizable across new sites (Wramneby et al., 2008). Model calibration and inclusion of parameter uncertainty via ensemble methods are rapidly becoming much more common practice in ecosystem modeling (Fischer et al., 2019; Fisher et al., 2019; Fer et al., 2018; Reichstein et al., 2019; Raczka et al., 2018). We informed our parameter distributions with an independent meta-analysis, but did not perform an additional calibration, and this decision may have increased our estimated process uncertainty. However, the overall contribution of parameter uncertainty was relatively modest (18%) suggesting parameter variance is not a dominant source of uncertainty in our analysis.

We found that meteorological uncertainty was increasingly important over decadal hindcasts (Figure 7). While hindcasting in this circumstance is much easier than forecasting because we hindsight allows us to know what happened over the last 50 years at Harvard Forest, we deliberately chose meteorological drivers to mimic the uncertainty of driver uncertainty in a true forecast. Our work agrees with studies showing the importance of carefully constructing future climate scenarios, as climate uncertainty increasingly contributes to total forecast variance (Feddema et al., 2005). Our analysis adds to previous work suggesting that meteorological uncertainty increases over time in comparison to parameter uncertainty (Lovenduski and Bonan, 2017; Bonan et al., 2019) by showing that the covariance between initial condition uncertainty and meteorological uncertainty may be contributing to the long-term increase of total forecast uncertainty observed by more complex forest demographic models (Raczka et al., 2018). Constraining the starting conditions of a forest stand gave us much more predictive power on the effects of varying climate on future stand productivity.

### 4.2 Initial conditions

Initial conditions have long been shown to affect successional pathways in temperate forests (Myster and Pickett, 1990). We found that initial condition uncertainty was the dominant source of uncertainty over our 60 year hindcast (Figure 7, hashed areas). Our findings on initial conditions agree with Alexander et al. (2017) and Ge et al. (2018) showing that model spin-up, and underlying equilibrium assumptions, could lead to very large, persistent uncertainties (Figure 5 left versus right). While it may be difficult to find field data to derive initial condition uncertainty estimates for longer term model simulations, it is always possible to construct informative priors about initial condition states from past ecological literature (Cressie et al., 2009; Hobbs and Hooten, 2015). Our analysis suggests that focusing efforts on data constrained initialization will be the most successful approach for improving forecast accuracy across forest gap models (68% reduction in total forecast variance from data constraints), even on multi-decadal timescales.

Large scale data are becoming increasingly available for terrestrial ecosystem model initialization, with advances in airborne and remote sensing measurements being particularly transformative. Not only can optical measurements be used to map canopy properties like leaf area index (LAI), but recent technological advancements in lidar, radar, and microwave remote sensing have improved our ability to map structural plant characteristics, like volume and canopy height, that are more directly related to terrestrial carbon pools, like total aboveground biomass (Goetz et al. 2009; Le Toan et al. 2011; Chave et al. 2019; Schepaschenko et al. 2019; Smith et al. 2020) as well as abiotic initial conditions, such as soil moisture (Entekhabi et al., 2010). In addition, hyperspectral remote sensing can provide us with the ability to map Plant Functional Type (PFTs) distribution globally (Shiklomanov et al., 2019). The Ecosystem Demography (ED) modeling team has demonstrated the effectiveness of remotely sensed Light Detection and Ranging (LiDAR) data for constraining initial conditions and decreasing near term forecast uncertainty (Hurtt et al. 2004; Antonarakis et al. 2014), a capacity that is now becoming applicable anywhere via the the Global Ecosystem Dynamics Investigation (GEDI) global LiDAR data product (Dubayah et al., 2020). These new measurements will provide terrestrial ecosystem models with significantly better data derived initial condition constraints than current spin up approaches (Schimel et al. 2015) and will greatly reduce uncertainty in both near-term and long-term forecasts of forested ecosystems.

### 4.3 Process uncertainty

Complex ecological systems often feature high dimensional interactions between state variables. This leads to process-based models that are highly and increasingly complex, which nonetheless remain imperfect representations of true ecosystem processes. In simpler ecological models, accounting for process uncertainty can result in more accurate predictions of modeled states as well as lead to ecological insights about which model processes need the most improvement (Cressie et al., 2009; Wikle, 2003). It is unclear if the trend towards increased model complexity in forest ecosystem models leads to increased predictive accuracy (Green et al., 2005; Hooten and Hobbs, 2015), as robust estimates of process uncertainty have, until now, been unavailable. The approach developed here moves beyond typical calculations of residual and validation errors and provides an estimate of process uncertainty that is dynamic (time-point to time-point) and quantifies the uncertainty remaining after accounting for all the other uncertainties discussed above, including observation error. Our approach allows us to propagate process uncertainty into ecological forecasts, which heretofore has generally been absent from process modeling approaches.

We found that process uncertainty contributes substantially to total forecast variance (Figure 7 and 9, pink). The particular process model we used, LINKAGES version 1.0, is something of a classic (Bonan et al., 2002), but it is an older model that has since been replaced in most modeling applications. It may be that an alternative process model would be better at predicting 60 years of forest stand dynamics that still requires testing to prove. We do note that the process covariance estimation itself is quite small (Table 3), suggesting that LINKAGES does adequately capture annual forest development changes. However, a small annual error is magnified over time, resulting in large 50 year uncertainty. Our estimate of process error covariance in LINKAGES over the data assimilation time period (50 years) suggests there are errors in modeled species mortality and recruitment, especially in red maple, that led to notable process uncertainty over many years. This finding aligns with previous studies in New England showing that the competitive relationships between red maple and red oak are difficult to understand and therefore predict (Lorimer, 1984; Abrams, 1998; Eisen and Plotkin, 2015).

The large impact of process uncertainty on forecast certainty suggests that future efforts might benefit from parsing out specific components of process uncertainty. This could include comparing alternative error models or the spatial and temporal autocorrelation in the process uncertainty, looking for evidence for heteroskedasticity, and partitioning of process uncertainty into bias and variance components. Some of the variability currently attributed to process uncertainty might also represent random individual variability (Clark et al., 2007) not currently captured by the model. Hierarchical modeling approaches (Clark et al., 2005) provide a means of partitioning this variability (Dietze, 2017b). New emulator methods are emerging to apply hierarchical approaches to complex process models (Fer et al., 2018).

### 4.4 Soil carbon

Constraining belowground soil carbon with aboveground productivity inputs has been a hallmark of our understanding of the evolution of long-term soil carbon accumulation (Meentemeyer, 1978; Aber, 1982; Solomon, 1986). LINKAGES mechanistically links aboveground biomass production, which was well constrained in our model thanks to SDA, to soil carbon through input from litter and tree mortality. But, the pools of soil carbon were not constrained by aboveground inputs in our model and grew to unrealistic levels. Variance partitioning reveals that this lack of constraint is caused by both process and initial condition uncertainty (Figure 9). But, parameter and meteorological uncertainty also yield hindcasts that are far from reality (Figure 8). While the link between aboveground and belowground pools has typically been assumed to be a quadratic cumulative relationship (Jenny, 1941), our work suggests that more evaluation is necessary to determine a better modeled representation. We also must add that, despite diligent model testing, it is possible that errors in our version of LINKAGES might have produced this result. Some forest gap models have alternative links between aboveground inputs and belowground pools (Friend et al., 1997), but it is unclear if more complex processes or different processes would reduce forecast uncertainty. In order to improve the link between aboveground inputs and belowground accumulation we agree with the sentiment in (Huber et al., 2020) that multiple model representations of unclear mechanistic processes should be used for predictions. We suggest that future directions focus on incorporating a variety processes known to affect the evolution of soil carbon beyond aboveground inputs using ensemble based methods. Furthermore, more variance partitioning exercises like those demonstrated here would efficiently point to which aspects of soil process modeling need the most attention in order to forecast long-term soil carbon.

### 4.5 Future Directions

Beyond forest gap models, our results call into question the conventional wisdom in many areas of ecological modeling more broadly, such as the reliance on “spin-up” initial conditions and the exclusion of process uncertainty from predictions. Most ecological forecasts are made without the inclusion of key uncertainties, with many made purely deterministically, leaving out uncertainty quantification altogether (Cressie et al., 2009). This common practice creates projections that may be precise but are often inaccurate. Similarly, many ecological fields focus on specific aspects of uncertainty without considering the full suite of possible sources of uncertainty. In particular, a lesson we learned, that demographic stochasticity is not the dominant uncertainty on forest gap models, likely extends more broadly, suggesting that the current reliance on specific uncertainties or stochasticities in other ecosystem modeling fields may be misleading ecologists about the dominant drivers of uncertainty. Similarly, many ecological projections have focused on uncertainty in parameters and meteorological drivers (Kremer, 1983; Eberhardt, 1987; Regan et al., 2002; Grimm et al., 2005; Zwart et al., 2019). While it is clear that these uncertainties do contribute to ecological modeling in general, it remains unclear what the relative contributions of parameter and meteorological uncertainty are to total forecast uncertainty across different spatial and temporal scales. For example, conventional wisdom suggests that initial condition uncertainties are likely to decrease over time, and climate scenario uncertainty is likely to increase with time (Cox and Stephenson, 2007; Dietze, 2017b). This crossover could vary enormously across systems, as could the impacts of other uncertainties.

Improving predictions of ecosystems properties not only leads to more efficient progress in advancing basic research and theory but also enhancements for decision makers and stakeholders. We demonstrated the power that uncertainty quantification has in ecology to reveal which long-standing modeling assumptions (spin-up initial conditions are sufficient; models with demographic stochasticity included are sufficient to capture uncertainties; process uncertainty is negligible) are not upheld by data and what steps can be taken to immediately increase forecast accuracy in forest gap modeling. These lessons are not unique to forest ecology. Moving forward, there is a critical need to extend analyses like these to more ecosystems, additional models, and larger spatial and temporal scales. This extension will allow ecologists to assess the generalities of our conclusions and to understand variation of the relative importance of different uncertainties across systems (Dietze, 2017b). Demographic stochasticity, parameter, meteorological, and process uncertainties are quantities that are measured across scales and systems. These types of methodologies can be used to quantitatively move toward better and more useful ecological predictions through systematic evaluation of the contribution of each uncertainty to total forecast uncertainty across different scales and systems.

## Acknowledgements

This material is based upon work supported by the National Science Foundation MacroSystems Biology under grant no. DEB-1241874, 1241891, 1458021. The authors thank the PalEON and PEcAn teams for support in early development of this project. We also thank Harvard Forest scientists Neil Pederson and Audrey Barker-Plotkin for collecting and archiving the field data behind the aboveground biomass estimates.

## Author Contributions

AR, MD, and JM designed project; AR and MD developed the data assimilation algorithm; AR analyzed data with advice from MD and JM; AR, JM, and MD wrote paper with further text contributions from all authors in the Supplementary Materials. Specifically, AD created and fit the tree ring model to create the species level biomass estimates, CR downscaled the climate drivers used to drive the model in this analysis, and JT fit the model that calculated the climate driver weights. All data and code used for this analysis is available in the manuscript supplementary materials or publicly on GitHub.

## 5 Data Availability Statement

All data for analysis, analysis workflow scripts, and post processing scripts can be found at PEcAn workflow IDs 1000009667, 1000009397, and 1000010510. ID 1000009667 contains the bulk of the variance partitioning output. ID 1000010510 and 1000009397 contain the model runs for understanding the effects of updating the parameters with prior information. Furthermore, the analysis scripts are available as part of the assim.sequential package within PEcAn at: https://github.com/PecanProject/pecan/tree/develop/modules/assim.sequential.

