## Supplemental Document for "Towards understanding predictability in ecology: A forest gap model case study"

### Supplemental Materials

#### 1 Meteorological Climate Ensemble Weights

##### 1.1 Statistical filtering of downscaled general circulation climate model output

In this section, we outline the procedure to filter the downscaled general circulation model (GCM) output to be consistent with gridded statistical paleoclimate reconstructions. The climate products that we use for filtering are the Mann et al. (2009) reconstruction of mean annual temperature and the Living Blended Drought Atlas (LBDA) from Cook et al. (2010) for mean annual Palmer Drought Severity Index (PDSI). The Mann et al. (2009) data product contains mean temperature estimates starting in AD 500 and ending in AD 2005 at a grid-cell resolution of $5^\circ \times 5^\circ$ covering much of the globe. The Cook et al. (2010) data product contains mean annual PDSI estimates starting in AD 0 and ending in AD 2006 at a grid cell resolution of $0.5^\circ \times 0.5^\circ$ covering much of North America.

##### 1.2 Model

The goal of the analysis is to produce a vector of weights for each member of the climate model ensemble for each year in the period 0 AD - 2000 AD. Let the empirical observations for year $t$ be $\mathbf{y}_t = (T_t, PDSI_t)'$ where $T_t$ is the Mann mean temperature estimate and $PDSI_t$ is the LBDA mean estimate at the grid cell of interest for year $t$. For the $i$th climate model run out of an ensemble of $N$ different climate runs, let $\mathbf{x}_{it} = \left(\tilde{T}_{it}, \widetilde{PDSI}_{it}\right)'$ be the climate model estimate of mean annual temperature and mean annual PDSI for the grid cell of interest for year $t$. Then, the goal is to find weights $\mathbf{w}_t = (w_{1t}, \ldots, w_{Nt})'$ that minimize the squared difference

$$\underset{\mathbf{w}_t}{\text{argmin}} \left\| \mathbf{y}_t - \sum_{i=1}^{N} w_{it}\mathbf{x}_{it} \right\|^2, \tag{1}$$

subject to the constraint that $\sum_{i=1}^{N} w_{it} = 1$. A primary goal for performing a statistical data assimilation is the preservation of variability in the weighted climate model ensemble. Model (1) has a significant issue in that the weight vector $\mathbf{w}_t$ could place very large weight on a single realization or a small handful of realizations. This is especially likely if the empirically estimated climate state is not close to the climate model ensemble. If this occurs, there will be an undesirable reduction in the variability of the weighted climate model ensemble. One way to preserve the variability in the weighted climate model ensemble is to bootstrap the empirical climate realizations using predictive standard deviations. For example, let $\mathbf{y}_t^{(k)}$ be the $k$th bootstrap draw from the predictive distribution $\left[\mathbf{y}_t^{(k)} \middle| \mathbf{y}_t, \mathbf{\Sigma}_{y_t}\right]$, where $\mathbf{\Sigma}_{y_t} = \text{diag}\left(\sigma^2_{T_t}, \sigma^2_{PDSI_t}\right)$ is the predictive covariance that is assumed to be diagonal and is estimated by comparing the reconstruction predictions to the observation-era records during the modern period. Ideally, the predictive standard deviations would be included in the data product and would increase further back in time as the amount of empirical proxy data used in the reconstruction products decreases; however, the predictive standard deviations are not reported as part of the data products used. Despite this, the predictive standard deviations can be approximated by comparing the output of modern-era data products to the Mann et al. (2009) and Cook et al. (2010) empirical paleoclimate proxy products. The modern-era climate proxies used in estimating the predictive standard deviations are the parameter-elevation regression on independent slopes model (PRISM Climate Group, Oregon State University 2004) product for mean annual temperature and the annual mean PDSI data product from Dai, Trenberth, and Qian (2004)[1] where we extract the record at the location of interest during the period of overlap in the modern era and calculate the root mean sum of squares as an estimate of the predictive standard deviation. In truth, one would expect these uncertainties to increase further in the past as paleoclimate data used to fit the models becomes more sparser; however, we treat these as constant in time as a first order approximation. Then, applying Eq. (1) to each of the bootstrap samples (assuming a Gaussian predictive distribution) produces a bootstrap distribution of weights $\mathbf{w}_t^{(k)}$ and the final weighted model ensemble is given by the bootstrap-averaged weights

$$\bar{\mathbf{w}}_t = \frac{1}{K} \sum_{k=1}^{K} \underset{\mathbf{w}_t^{(k)}}{\text{argmin}} \left\| \mathbf{y}_t^{(k)} - \sum_{i=1}^{N} w_{it}^{(k)} \mathbf{x}_{it} \right\|^2 . \tag{2}$$

Thus, the bootstrap-averaged weights show less degeneracy and do a better job of preserving the variability in the weighted model ensemble.

### 2 Prior Distributions

There are 21 parameters per species in LINKAGES. We used Harvard Forest field measurements to improve parameter representation and incorporate parameter uncertainty in the following PEcAn workflow (LeBauer et al., 2013). First, we defined Bayesian prior distributions for each parameter. We began by giving each parameter an uninformative prior distribution with a mean based on the default LINKAGES parameterization. To identify the parameters most needing constraint and reduce the dimensionality of subsequent

[1]Dai Palmer Drought Severity Index data provided by the NOAA/OAR/ESRL PSD, Boulder, Colorado, USA, from their Web site at `https://www.esrl.noaa.gov/psd/`

parameterization efforts, we conducted a one-at-a-time parameter uncertainty analysis. Using a Bayesian meta-analysis (LeBauer et al., 2013), we were able to constrain specific leaf area (SLA) for each species with trait data from the (BETY) database (LeBauer et al., 2018), and allometric and recruitment parameters with information from Catovsky and Bazzaz (2000), Dietze et al. (2008), and Sullivan et al. (2017).

#### 2.1 Red Maple Prior Distributions

| | distn | parama | paramb | n |
|---|---|---|---|---|
| SLA | weibull | 0.50 | 10.00 | 2548 |
| DMAX | norm | 6600.00 | 1.00 | |
| DMIN | norm | 1000.00 | 1.00 | |
| AGEMX | unif | 100.00 | 200.00 | |
| SPRTND | norm | 13.00 | 1.00 | |
| SPRTMX | norm | 200.00 | 1.00 | |
| D3 | unif | 0.25 | 0.75 | |
| CM1 | unif | 2.69 | 2.89 | |
| CM2 | norm | 219.77 | 1.00 | |
| CM3 | unif | 0.00 | 0.00 | |
| CM4 | norm | -0.60 | 0.10 | |
| CM5 | norm | 0.85 | 0.10 | |
| FWT | unif | 400.00 | 480.00 | |
| Gmax | norm | 100.00 | 1.00 | |
| MPLANT | lnorm | 4.45 | 1.00 | |
| FROST | norm | -17.00 | 2.00 | |
| SLTA | norm | 0.01 | 0.25 | |
| SLTB | norm | 0.37 | 0.25 | |
| HTMAX | norm | 15.00 | 1.00 | |
| DBHMAX | norm | 0.86 | 0.03 | |

Table 1: Red Maple Prior Distributions

#### 2.2 Yellow Birch Prior Distributions

| | distn | parama | paramb | n |
|---:|---|---:|---:|---:|
| SLA | weibull | 0.50 | 10.00 | 2548 |
| DMIN | norm | 1100.00 | 1.00 | |
| HTMAX | norm | 30.00 | 1.00 | |
| DBHMAX | norm | 0.80 | 0.03 | |
| AGEMX | unif | 150.00 | 350.00 | |
| Gmax | norm | 50.00 | 1.00 | |
| SPRTND | norm | 43.00 | 1.00 | |
| SPRTMX | norm | 100.00 | 1.00 | |
| SPRTMN | norm | 12.00 | 1.00 | |
| D3 | unif | 0.25 | 0.75 | |
| FWT | unif | 200.00 | 300.00 | |
| CM1 | unif | 2.84 | 3.04 | |
| CM2 | norm | 117.52 | 1.00 | |
| CM3 | unif | 0.00 | 0.00 | |
| CM4 | norm | -1.20 | 0.10 | |
| CM5 | norm | 1.30 | 0.10 | |
| DMAX | norm | 2500.00 | 1.00 | |
| MPLANT | lnorm | 6.83 | 1.00 | |
| FROST | norm | -18.00 | 2.00 | |
| SLTA | norm | 0.07 | 0.25 | |
| SLTB | norm | 0.01 | 0.25 | |

Table 2: Yellow Birch Prior Distributions

#### 2.3 Beech Prior Distributions

| | distn | parama | paramb | n |
|---|---|---|---|---|
| SLA | weibull | 0.50 | 10.00 | 2548 |
| DMIN | norm | 1326.00 | 50.00 | |
| AGEMX | weibull | 2.00 | 494.97 | |
| Gmax | unif | 52.00 | 92.00 | |
| SPRTND | norm | 172.00 | 1.00 | |
| SPRTMX | norm | 30.00 | 1.00 | |
| MPLANT | unif | 0.00 | 60.00 | |
| HTMAX | norm | 30.00 | 1.00 | |
| DBHMAX | lnorm | 1.00 | 1.00 | |
| D3 | unif | 0.25 | 0.75 | |
| FWT | unif | 400.00 | 480.00 | |
| CM1 | unif | 2.84 | 3.04 | |
| CM2 | norm | 117.52 | 1.00 | |
| CM3 | unif | 0.00 | 0.00 | |
| CM4 | norm | -1.20 | 0.10 | |
| CM5 | norm | 1.30 | 0.10 | |
| root2shoot | beta | 1.00 | 1.00 | |
| DMAX | norm | 5537.00 | 10.00 | |
| FROST | unif | -100.00 | 0.00 | |
| SPRTMN | norm | 6.00 | 1.00 | |
| SLTA | norm | 0.57 | 0.25 | |

Table 3: Beech Prior Distributions

#### 2.4 Red Oak Prior Distributions

| | distn | parama | paramb | n |
|---:|---|---:|---:|---:|
| SLA | weibull | 0.50 | 10.00 | 2548 |
| D3 | unif | 0.25 | 0.75 | |
| FWT | unif | 400.00 | 480.00 | |
| CM1 | unif | 2.84 | 3.04 | |
| CM2 | norm | 117.52 | 1.00 | |
| CM3 | unif | 0.00 | 0.00 | |
| CM4 | norm | -1.20 | 0.10 | |
| CM5 | norm | 1.30 | 0.10 | |
| DMAX | norm | 4571.00 | 1.00 | |
| DMIN | norm | 1100.00 | 1.00 | |
| AGEMX | unif | 300.00 | 500.00 | |
| Gmax | norm | 130.00 | 1.00 | |
| SPRTND | norm | 32.00 | 1.00 | |
| SPRTMX | norm | 40.00 | 1.00 | |
| SPRTMN | norm | 17.00 | 1.00 | |
| MPLANT | lnorm | 2.64 | 1.00 | |
| FROST | norm | -18.00 | 2.00 | |
| SLTA | norm | -0.30 | 0.25 | |
| SLTB | norm | 0.52 | 0.25 | |
| HTMAX | norm | 48.00 | 1.00 | |
| DBHMAX | norm | 1.50 | 0.03 | |

Table 4: Red Oak Prior Distributions

#### 2.5 Hemlock Prior Distributions

| | distn | parama | paramb | n |
|---|---|---|---|---|
| SLA | weibull | 0.50 | 10.00 | 2548 |
| DMIN | norm | 1324.00 | 10.00 | |
| HTMAX | unif | 30.00 | 40.00 | |
| DBHMAX | unif | 1.00 | 2.00 | |
| AGEMX | weibull | 2.00 | 1060.66 | |
| Gmax | unif | 27.00 | 67.00 | |
| D3 | unif | 0.25 | 0.75 | |
| CM1 | unif | 2.69 | 2.89 | |
| CM2 | norm | 219.77 | 1.00 | |
| CM3 | unif | 0.00 | 0.00 | |
| CM4 | norm | -0.60 | 0.10 | |
| CM5 | norm | 0.85 | 0.10 | |
| FWT | unif | 400.00 | 480.00 | |
| root2shoot | beta | 1.00 | 1.00 | |
| DMAX | norm | 3800.00 | 10.00 | |
| MPLANT | lnorm | 6.58 | 1.00 | |
| FROST | norm | -13.00 | 2.00 | |
| SLTA | norm | -0.02 | 0.25 | |
| SLTB | norm | 0.36 | 0.25 | |

Table 5: Hemlock Prior Distributions

### 3 Ecosystem Model Description: LINKAGES

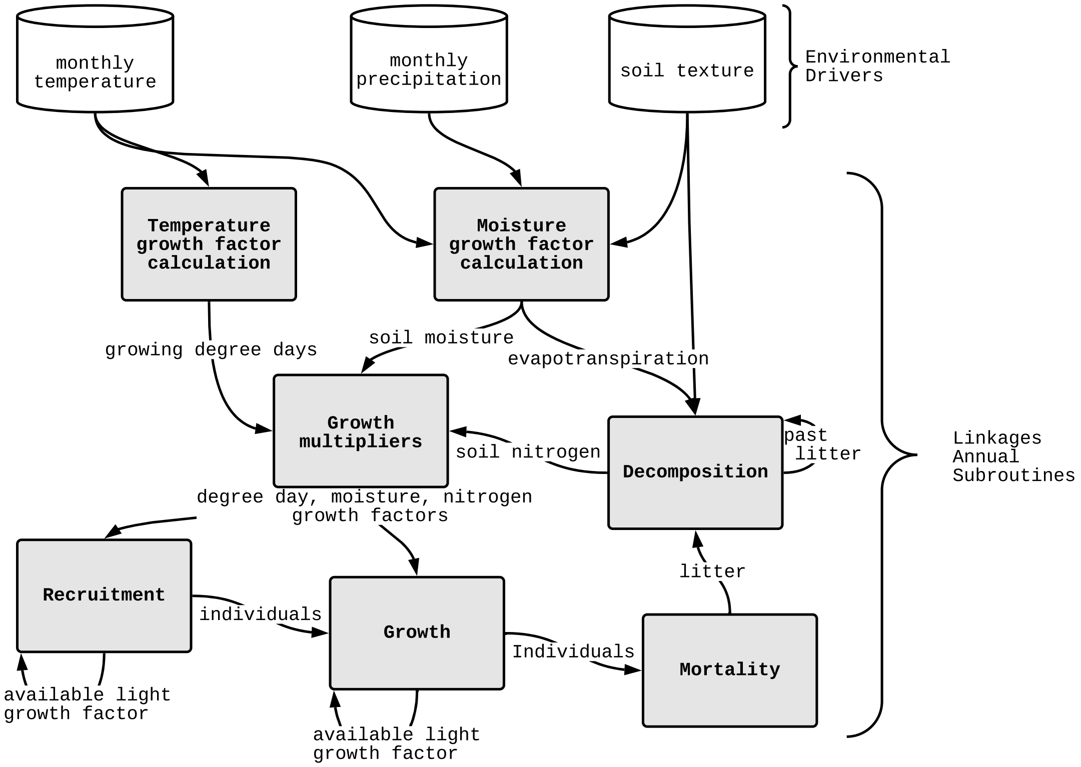


Figure 1: Conceptual diagram of the forest gap model LINKAGES (Post and Pastor 1996). Environmental driver data is used by subroutine temperature, moisture, and decomposition to calculate annual growth factors for growing degree day, soil moisture, and soil nitrogen in the growth multipliers subroutine. Growth multipliers subroutine calculates species specific growth factors using species specific parameters (description of parameters available in supplemental materials figure 2). These annual growth factors along with diameter at breast height (DBH), age, and growing status of each individual are input into recruitment, growth, and mortality subroutines. Within the recruitment, growth, and mortality subroutines individual DBH, age, and growing status are incremented and individuals can be recruited or be removed (mortality) based on site conditions. Available light growth multipliers are calculated for each individual during the growth subroutine. Litter pools are saved year to year and contribute to available nitrogen and soil nitrogen growth factors calculated in the decomposition subroutine.

#### 3.1 Parameter Uncertainty Analysis Results

We also found that climate related parameters related to growing degree day minimum and maximum and frost tolerance were causing LINKAGES to consider some species not viable for growth. For these climate envelope parameters, we expanded the priors to include values within the range of our meteorological data centered on the LINKAGES default parameter values. We found that parameters affecting total aboveground biomass sensitivity varied between species (Figure 2). Red maple biomass is most sensitive to nitrogen growth multiplier 1 (CM1) and somewhat sensitive to maximum seeding rate per plant (MPLANT). Yellow birch and beech are most sensitive to number of sprouts per stump (SPTRND). Red oak has the most sensitive to nitrogen growth multiplier 2 (CM2).

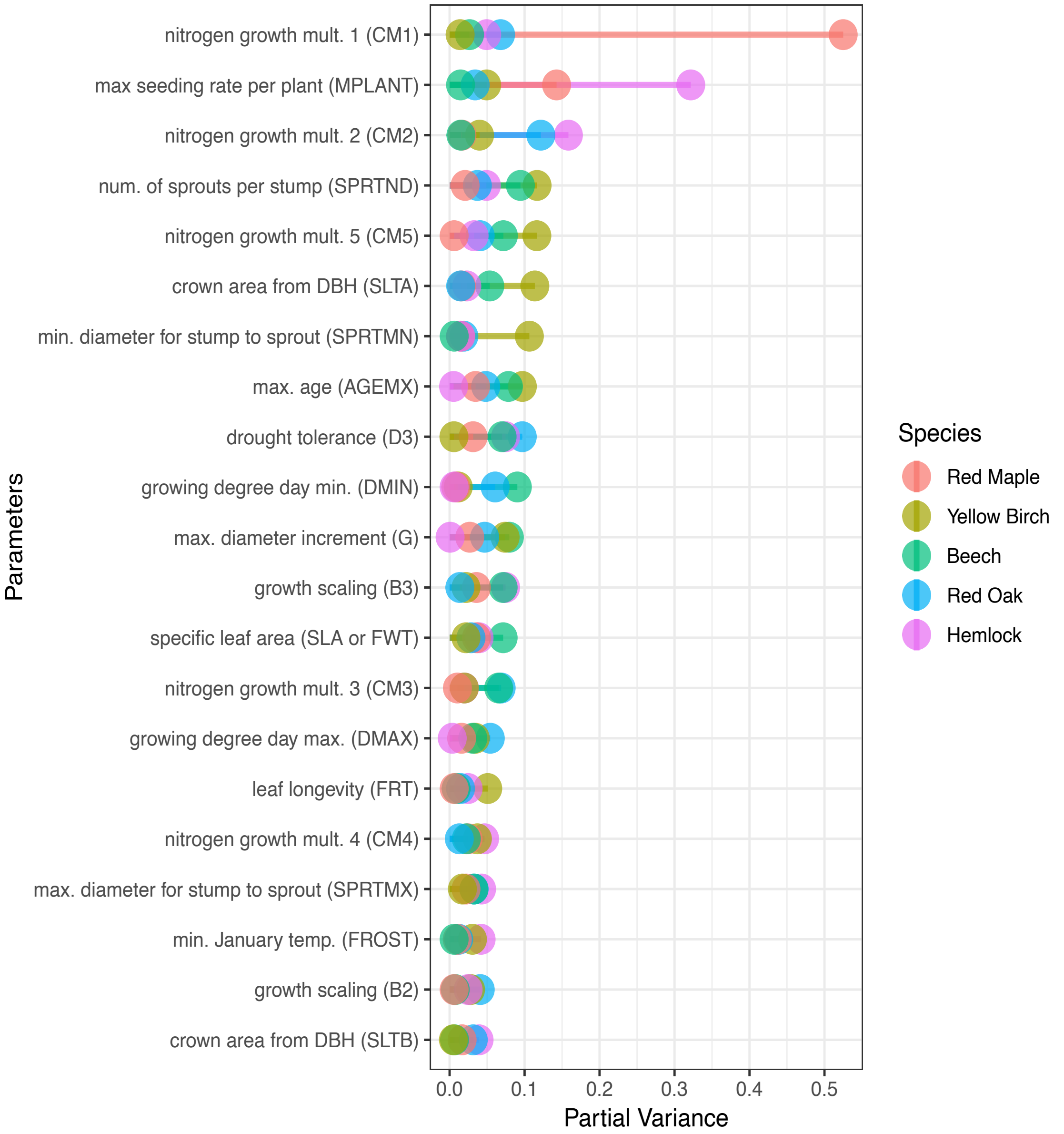


Figure 2: Partial variance of LINKAGES 21 species specific parameters calculated from full sensitivity analysis on output variable total aboveground biomass.

### 4 Statistical Description of the Tobit Wishart Ensemble Filter (TWEnF)

Before performing each analysis step, we first have to calculate the forecast mean and covariance. In the traditional EnKF, this is the ensemble sample mean ($\boldsymbol{\mu}_f$) and covariance ($\boldsymbol{P}_f$), but because we applied the Tobit transformation, we estimated $\boldsymbol{\mu}_f$ and $\boldsymbol{P}_f$ with a Normal-inverse-Wishart as follows

$$
\begin{aligned}
z_{ij} &= \begin{cases} 0 & x_{ij} \leq y_L \\ 1 & x_{ij} > y_L \end{cases} \\
\boldsymbol{x}_i &\sim \mathrm{N}(\boldsymbol{\mu}_f, \boldsymbol{P}_f)^{w_i} \\
\boldsymbol{\mu}_f &\sim \mathrm{N}(\boldsymbol{\mu}_0, \boldsymbol{\Sigma}_0) \\
\boldsymbol{P}_f &\sim \mathrm{Inv-Wishart}(\boldsymbol{S}, \nu),
\end{aligned}
$$

where $\boldsymbol{\Sigma}_0$ and $S$ are prior covariances (on the forecast mean and covariance respectively), $\nu$ is the number of elements in the forecast mean vector ($\boldsymbol{\mu}_f$), $\boldsymbol{\mu}_0$ is prior mean vector, $\boldsymbol{x}_i$ is a vector of predicted state variables for each forecast ensemble member $i$, $z_{ij}$ is an indicator variable indicating which state variable ($j$) of each ensemble member ($i$) is outside of the truncated space, and $y_L$ is the left truncation for the Tobit, which equals 0 in our case. Finally, $w_i$ is the known weight associated with each of the ensemble members based on an *a priori* analysis of the meteorological drivers described in the supplemental materials section 1.1. We fit this model in the R version 3.4.4 computing environment using the 'nimble' package version 0.8.0 (Team et al. 2017; de Valpine et al. 2017) assuming vague priors. We used a user defined 'toggle' sampler that samples $\boldsymbol{x}_i$ with a random walk sampler when $z_{ij} = 0$. We also wrote a user defined conjugate sampler for $\boldsymbol{P}_f$ to incorporate ensemble weights $w_i$. These are available in the PEcAn GitHub repository (`https://github.com/PecanProject/pecan`).

With $\boldsymbol{\mu}_f$ and $\boldsymbol{P}_f$ now in hand, the TWEnF analysis step is based off the EnKF, but in the TWEnF we added a latent state vector ($\boldsymbol{x}_l$) that accounts for added process covariance and non-normal errors. Like the previous step, we begin with a Tobit transformed likelihood (Eqn. 1), but this time we are using the Tobit to account for zero truncation and zero inflation in the data, rather than the forecast. The following TWEnF was fit at every time step.

$$
\begin{aligned}
y_j &= \begin{cases} y_j^* & \text{if } y_j^* > y_L \\ y_L & \text{if } y_j^* \leq y_L \end{cases} && (3) \\
\boldsymbol{y}^* &\sim \mathrm{MVN}(\boldsymbol{x}_l, \boldsymbol{R}) && (4) \\
\boldsymbol{x}_l &\sim \mathrm{MVN}(\boldsymbol{x}_f, \boldsymbol{Q}) && (5) \\
\boldsymbol{x}_f &\sim \mathrm{MVN}(\boldsymbol{\mu}_f, \boldsymbol{P}_f) && (6) \\
\boldsymbol{Q} &\sim \mathrm{Inv-Wishart}(\boldsymbol{\Omega}_q, \beta_q), && (7)
\end{aligned}
$$

where $\boldsymbol{y}^* = (y^*, ..., y_J^*)$ is the mean data vector calculated from the posterior of the tree ring aboveground biomass data product, $\boldsymbol{R}$ is the observation error covariance calculated from the posterior draws of the same data product, $\boldsymbol{x}_l$ is the latent model state that is estimated from the forecast ensemble mean $\boldsymbol{x}_f$ and process covariance $\boldsymbol{Q}$. Let $\boldsymbol{\Omega}_q$ and $\beta_q$ be the prior shape parameters for the process covariance that are estimated over time, as described in the next paragraph. This model was also fit in using 'nimble'. We used a random walk block sampler to fit $\boldsymbol{x}$ and $\boldsymbol{x}_f$ where there are data from $\boldsymbol{y}$ (Roberts and Sahu, 1997). We used slice samplers for $\boldsymbol{x}$ and $\boldsymbol{x}_f$ where there are no data from $\boldsymbol{y}$ (Neal, 2003). In our case, we also used a slice sampler for the soil carbon pool because we do not have data directly informing this state variable. We again used the toggle sampler for $\boldsymbol{y}$ that contain zeros and we used a conjugate (Gibbs) sampler for $\boldsymbol{Q}$.

From the MCMC posterior draws of $\boldsymbol{x}_l$, we calculate the analysis mean ($\boldsymbol{\mu}_a$) and covariance ($\boldsymbol{P}_a$). The analysis mean and covariance are then used, as they would be in a ensemble adjustment filter (Anderson, 2001), to "nudge" the forecast ensemble members to have the correct posterior mean and covariance. After the adjustment, we restarted each ecosystem model ensemble member with an updated state vector. The specific LINKAGES restart scripts are described in Supplemental Section 5. We updated the estimate of the process covariance ($\boldsymbol{Q}_t$) every time step by updating the shape parameters of the Inverse-Wishart distribution as follows

$$\Omega_{q_{t+1}} = \bar{Q}_t \beta_{q_{t+1}} \tag{8}$$

$$\beta_{q_{t+1}} = \mathrm{E}\left[\frac{\Omega_{rc}^2 + \Omega_{rr}\Omega_{cc}}{\mathrm{var}(\Omega_{rc})}\right]_t , \tag{9}$$

where $\Omega$ is the process precision to the process covariance $Q_t$ and $r$ and $c$ represent the rows and columns of the process precision matrix. In this step of the analysis, $\bar{Q}_t$ is the posterior mean estimate of $Q_t$ from the TWEnF. We assessed convergence at every time step using the Gelman-Rubin convergence diagnostic with the 'coda' package (Gelman et al., 1992; Plummer et al., 2006). Our workflow was restarted with different initial conditions if Gelman-Rubin diagnostics were greater than 1.01 for more than two monitored variables.

### 5 Ecosystem Model Restart

Once we have obtained the updated state variable matrix, we needed to restart our ecosystem model by rewriting the state variables into ecosystem model specific formats. For LINKAGES, we calculate new species density and individual tree DBHs so that the stand level biomass estimated in the MCEnF matches LINKAGES individual biomass calculations. To do this we first calculate each individual tree's forecast biomass and aggregate this to a species level. We determine a biomass correction factor by dividing the updated biomass from the MCEnF by the species biomass from the LINKAGES forecast. Then, we sample from the original pool of individuals based off the analysis biomass to obtain a new number of individuals. Adjusting the species level tree density first allows us to preserve stand age and size distribution from LINKAGES. Next, we adjust the individual's DBH directly using an optimization function to match the stand level biomass given from the MCEnF analysis. Soil carbon is much less complicated in LINKAGES, so we apply a soil carbon correction factor directly. These new model specific state variables are fed back into LINKAGES as current state variables.

### 6 Species Bias Diagnostics

By updating the state variables sequentially, data assimilation allows us to evaluate out five uncertainties each time the model was updated. We defined the model error, which is analogous to a residual, as the forecast minus the data. The update diagnostic, which shows the extent to which the model needs to be adjusted at each time step, is defined as the forecast minus the analysis (Dietze, 2017). Both of these diagnostics give us an indication of model bias. If the model bias increases or decreases over time, this indicates that the model process behind that state variable may need to be adjusted. For example, if the bias is increasingly positive for a specific species biomass, then we would suspect that the model is not representing mortality correctly for that species because the forecast is increasingly much larger than the analysis. Other types of more traditional diagnostics include root mean square error, correlation coefficient, and $R^2$ which are also used for testing the similarities and differences between the model forecast and the data (Hyndman and Koehler, 2006).

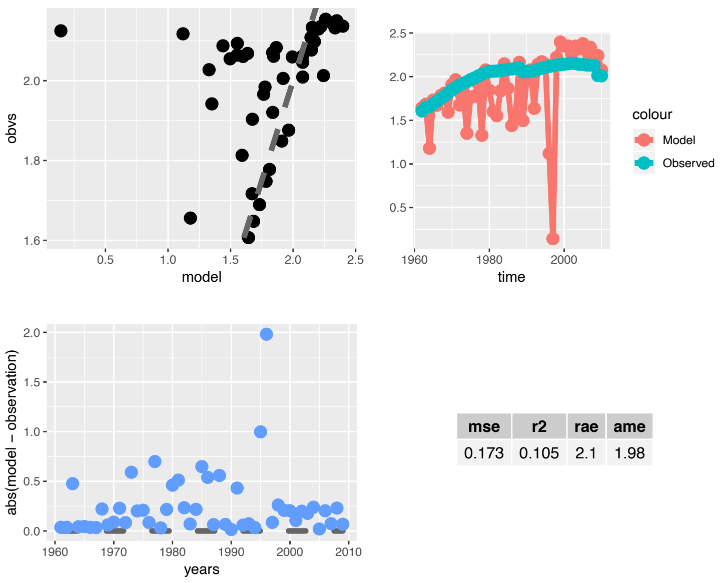


| mse | r2 | rae | ame |
|---|---|---|---|
| 0.173 | 0.105 | 2.1 | 1.98 |

Figure 3: Bias diagnostics for red maple

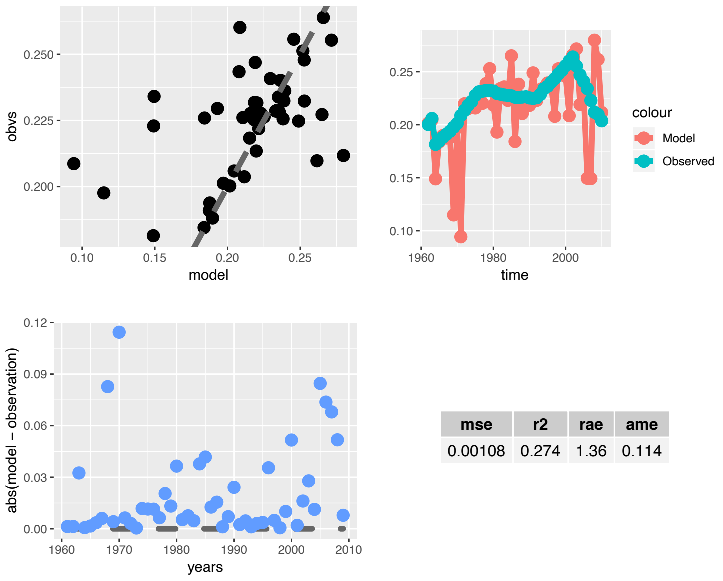


| mse | r2 | rae | ame |
|---|---|---|---|
| 0.00108 | 0.274 | 1.36 | 0.114 |

Figure 4: Bias diagnostics for yellow birch

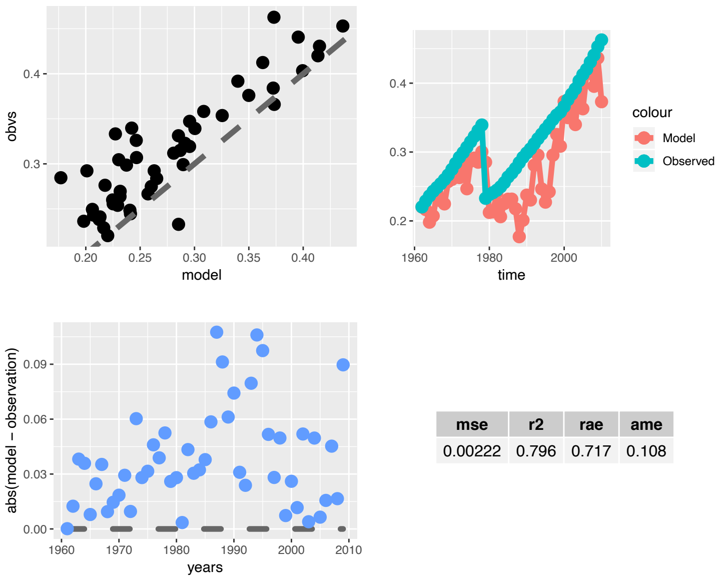


| mse | r2 | rae | ame |
|---|---|---|---|
| 0.00222 | 0.796 | 0.717 | 0.108 |

Figure 5: Bias diagnostics for American beech

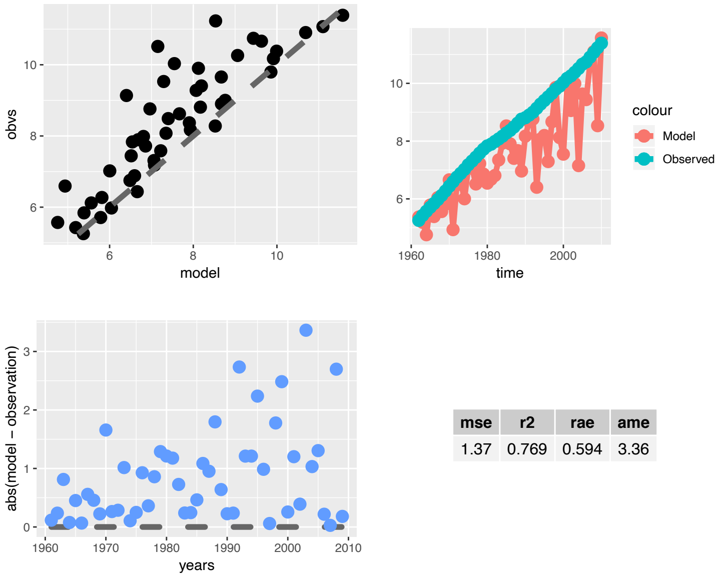


| mse | r2 | rae | ame |
|---|---|---|---|
| 1.37 | 0.769 | 0.594 | 3.36 |

Figure 6: Bias diagnostics for red oak

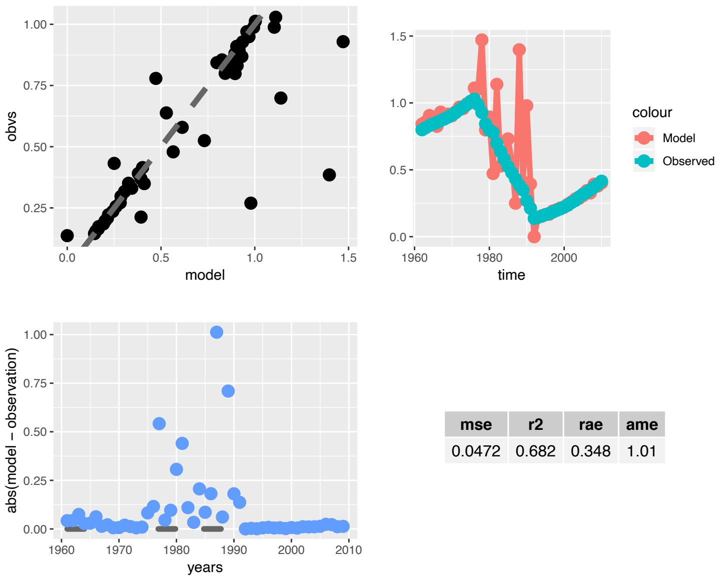


| mse | r2 | rae | ame |
|---|---|---|---|
| 0.0472 | 0.682 | 0.348 | 1.01 |

Figure 7: Bias diagnostics for eastern hemlock

### 7 Spin IC Species Level

Scenarios A1-4 were conducted with spin up initial conditions. In Supplemental Figure 8, we show the species level effects of the four other uncertainties. In contrast to the data-derived initial conditions scenarios (B1-4), these species-level uncertainties are not dominated by process uncertainty but also by parameter in red maple's case and meteorological for beech, red oak, and hemlock. This is because adding these uncertainties for 60 years before the analysis inflated the ensembles that continued inflated unconstrained by any data while the data constrained uncertainty scenarios did not reach saturation until the process uncertainty was included.

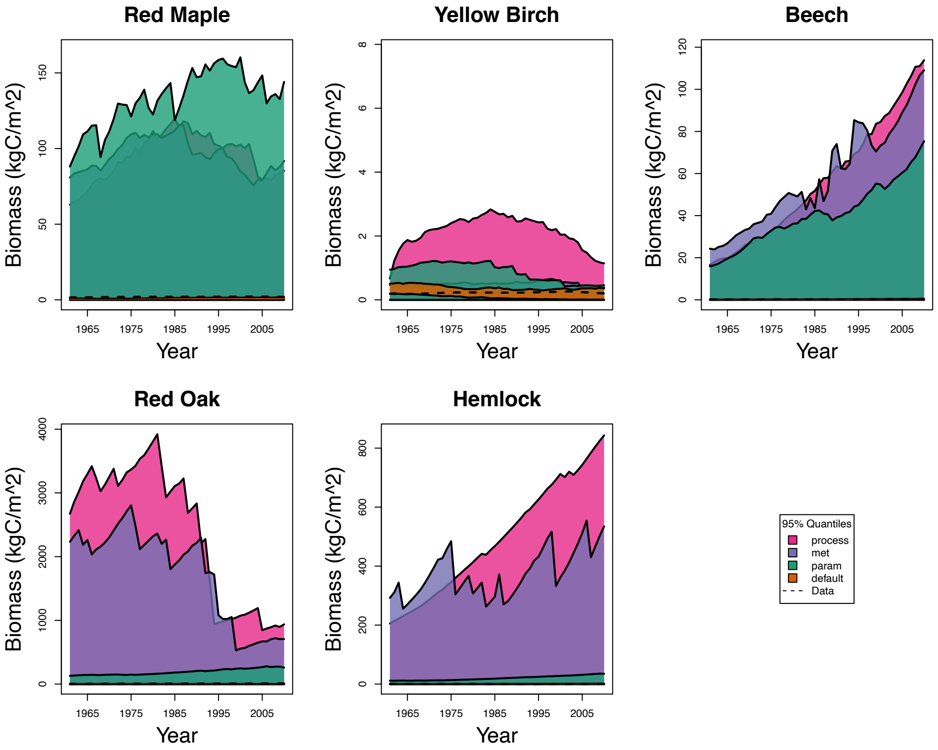


Figure 8: 95% quantiles of species level biomass over time colored by the four data spin-up initial condition uncertainty scenarios (scenarios A1-4). Variance from the first scenario arises only from demographic stochasticity (orange). Parameter, meteorological, and process uncertainty are added sequentially in the next three scenarios where the fourth scenario, with the widest credible intervals, contains variance from demographic stochasticity, parameters, meteorology, and process uncertainties. The dotted lines on all the plots are the 95% credible intervals of the data estimated from tree rings.

### 8 Forecast Covariance

Here we report the forecast covariance $P_f$ representing the covariance between species inherent in LINKAGES. Our estimation of process covariance $Q$ accounts for the covariance between species in the data that is beyond what is accounted for in the forecast or model covariance.

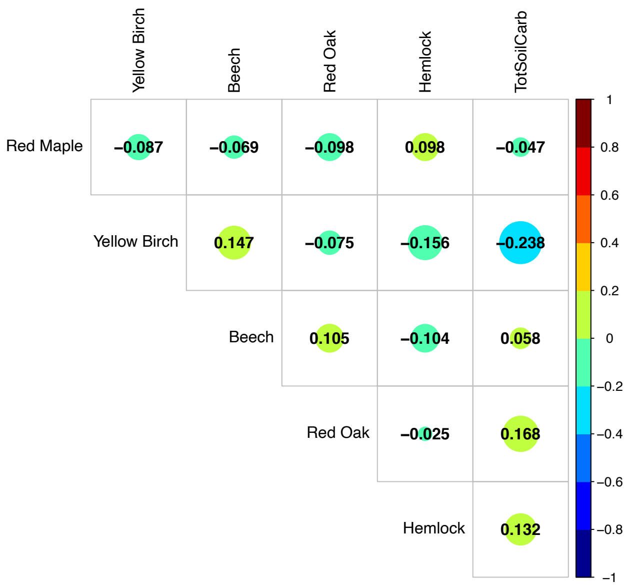


Figure 9: Correlations between species level biomass from the LINKAGES forecast.
